# Structural evolution of a yeast amyloid *in vivo* is shaped by chaperones

**DOI:** 10.64898/2026.08.10.743537

**Authors:** Ziang Wang, Samantha L. Weetman, Barbara Altenhuber, Alexey G. Murzin, James D. Gilbert, Riccardo Zenezini Chiozzi, Nadejda Koloteva-Levine, Helen R. Saibil, John Collinge, Mick F. Tuite, Wei-Feng Xue, Wenjuan Zhang

## Abstract

Cryo-electron microscopy (cryo-EM) studies of amyloid fibrils have revealed endpoint structures of disease-relevant filaments and polymorphic intermediates formed during *in vitro* assembly of prion-like proteins. However, how transmissible prion or prion-like amyloids evolve during *de novo* formation and maturation in living cells remains unknown. Here, using the yeast prion [*PSI^+^*] as a model, we isolated Sup35NM amyloid fibrils from successive stages of [*PSI^+^*] maturation in *Saccharomyces cerevisiae* and characterised their near-atomic structures and population-level structural diversity by combining cryo-EM and atomic force microscopy. We show that intermediate and mature states differ in predominant fibril structure and the regions of the Sup35 sequence incorporated into the core, and that structural diversity decreases during maturation. Curing of [PSI+] at the mature state by guanidine hydrochloride (GdnHCl), which selectively inhibits ATPase activity of the chaperone Hsp104, restored both the predominant intermediate amyloid structure and the broader structural diversity characteristic of the intermediate state. In addition, Hsp104, Ssa1 (Hsp70) and Sis1 (Hsp40) associate differently with fibrils from the two states. Together, these findings provide direct structural evidence for amyloid evolution *in vivo* and support a chaperone-mediated mechanism of conformer selection within a polymorphic amyloid population.

## Introduction

A prion is a proteinaceous infectious particle that lacks nucleic acid^1^. They form by undergoing a conformational shift from a normally soluble, monomeric protein into large, ordered, filamentous amyloid assemblies. Mammalian prions are infectious agents that cause fatal neurodegenerative diseases in both humans and animals and arise through structural transformation of the globular cellular prion protein (PrP^C^) into the misfolded, infectious amyloid conformation^2^. In other neurodegenerative diseases such as Alzheimer’s (AD) and Parkinson’s diseases and amyotrophic lateral sclerosis, it has been recognised that the pathological protein aggregates of tau, amyloid-β(Aβ), α-synuclein and TDP-43, may adopt a “prion-like” mechanism, where they spread and propagate through the nervous system in a similar way as PrP prions^3–5^. Molecular chaperones can fragment fibrillar assemblies into smaller seeds and thereby prompt the propagation of disease-related PrP^6^, tau^7^, and α-synuclein^8^ aggregates.

Recent advances in cryo-electron microscopy (cryo-EM) and image processing approaches for single-particle helical reconstruction by RELION^9^ have enabled near atomic-resolution structures of amyloid filaments from human post-mortem brains, revealing that specific folds in the cores of amyloid from tau, Aβ, α-synuclein, and TDP-43 define different neurodegenerative diseases^10^. These studies provided mechanistic insights into the structural basis of amyloid in diseases at the endpoint of protein assembly. More recently, time-resolved cryo-EM studies have captured structural snapshots during amyloid formation by characterising filament structures at successive time points during *in vitro* assembly of tau^11^ and human islet amyloid polypeptide (IAPP)^12^, revealing intermediate conformations and structural evolution during amyloid maturation. A recent study used a similar approach to reveal structural evolution of a functional amyloid that underlies age-related memory impairment in *Drosophila*^13^. However, the molecular mechanisms underlying *de novo* formation and maturation of prion or prion-like amyloids *in vivo*, and the roles of molecular chaperones in these processes, remain unknown.

The yeast prion [*PSI⁺*] arises from the conformational conversion of the translation termination factor Sup35^14–16^ protein in *Saccharomyces cerevisiae*. Sup35 amyloid in [*PSI⁺*] cells provides a tractable model for understanding amyloid formation and maturation *in vivo*. Sup35 contains an N-terminal prion-forming domain (N, residues 1-123), a solubilising and highly charged middle domain (M, residues 124-253), and a C-terminal domain essential for the translational termination function of Sup35 (residues 254-685)^17^. The first 114 residues are essential for the maintenance of [*PSI⁺*] phenotype^18^. The spontaneous *de novo* formation of [*PSI⁺*] is rare, but transient overexpression of the Sup35 or Sup35NM in yeast strains possessing [*PIN⁺*], another yeast prion formed by aggregation of the functionally uncharacterised protein Rnq1^19^, dramatically increases the rate of *de novo* [*PSI⁺*] induction^20^. During the *de novo* generation of [*PSI⁺*], an intermediate state in which Sup35 aggregates appear as ring-like shapes (hereafter termed the Ring state) has been typically observed before the aggregates transition into compact, dot-like puncta (hereafter termed the Dot state) characteristic of matured prions^21–24^. Cells containing Ring or Dot aggregates give rise to progeny with Ring and Dot aggregates, respectively, but Ring-containing cells frequently also generate progeny with diffuse fluorescence. Only Dot aggregates can transmit the [*PSI⁺*] phenotype to the mating cells^24^. Ring aggregates are bundles of long uninterrupted fibrils at the periphery of the cells, whereas Dot aggregates are short fibrils with diverse orientations in the cytoplasm^25,26^, reminiscent of observations from *in situ* cryo-ET data of Aβ amyloid in an AD human brain^27^.

Heat shock protein 104 (Hsp104) is a molecular chaperone and a powerful disaggregase in yeast. It is capable of disaggregating a wide range of protein aggregates and amyloid assemblies by utilising the energy derived from ATP hydrolysis^28,29^. The chaperone Hsp104, together with Ssa1 (Hsp70) and Sis1 (Hsp40), fragments Sup35 prion aggregates into smaller seeds that prompt propagation of [*PSI^+^*] and its transmission to daughter cells^30,31^. Hsp104 is essential for the propagation of almost all known yeast prions^30,32^. Cells can be cured of the [*PSI^+^*] prion by treatment with GdnHCl through its ability to selectively inhibit the ATPase activity of Hsp104^33,34^. Following GdnHCl treatment, the Dot state is reverted to the Ring state before complete prion clearance^21^. The structural basis of the intermediate Ring and mature Dot states of [*PSI⁺*] and the mechanisms by which chaperones influence the maturation and curing of [*PSI^+^*] remain poorly understood.

Here, we investigate *de novo* amyloid formation, structural evolution and maturation of prion particles *in vivo* using the yeast prion [*PSI^+^*] as a model. By combining AFM analysis of individual filaments with high-resolution cryo-EM, we quantified structural diversity within *ex vivo* Sup35NM amyloid populations and resolved the dominant Ring and Dot fibril structures at near-atomic resolution. We define structural diversity as the degree of variation among polymorphic structures within a fibril population. We further examined how molecular chaperones contribute to structural evolution of the Sup35 amyloid population during [*PSI^+^*] maturation.

## Results

### *De novo*-formed Sup35 aggregates at successive stages of [*PSI^+^*] maturation contain amyloid fibrils that can be isolated *ex vivo*

We overexpressed YFP-tagged Sup35NM in [*psi^−^*][*PIN^+^*] yeast cells to induce *de novo* formation of [*PSI^+^*]. These cells lack the N and M domains in their endogenous chromosomal SUP35 gene to avoid toxicity^24,26^. YFP or GFP tagging has been shown not to affect Sup35 assembly in yeast cells^21,26^. After 8 hours of Sup35NM overexpression, ribbon- and ring-like aggregates were readily detected in yeast cells by confocal microscopy. Small amounts of spot-like aggregates that have previously been proposed to represent oligomeric aggregates and/or phase separated droplets^35^, were also observed. Occasionally, large dot-like aggregates could be seen. After 24 h of Sup35NM overexpression, aggregates were detected in 52% of cells. Of these, 8% displayed large cytoplasmic puncta, whereas 92% showed ribbon- or ring-like aggregates at the cell periphery (**Fig. 1a**). These cells are hereafter referred to as Ring cells. To promote maturation of [*PSI^+^*] from the intermediate Ring state, Ring cells were re-streaked on plates for four times, corresponding to ∼100 generations. After this passage, only large dot-like puncta were observed inside these cells, hereafter referred to as Dot cells (**Fig. 1b**). Immunoblot analysis of sedimentation assays showed that more than 90% of Sup35 was present in insoluble aggregates in both Ring and Dot cells (**Extended Data Fig. 1a**). Co-localisation of both Ring and Dot aggregates with Amytracker and Thioflavin T (ThT) dyes that bind to fibrillar amyloid aggregates specifically confirmed that these aggregates contained amyloid fibrils (**Extended Data Fig. 1b**).

**Figure 1.**
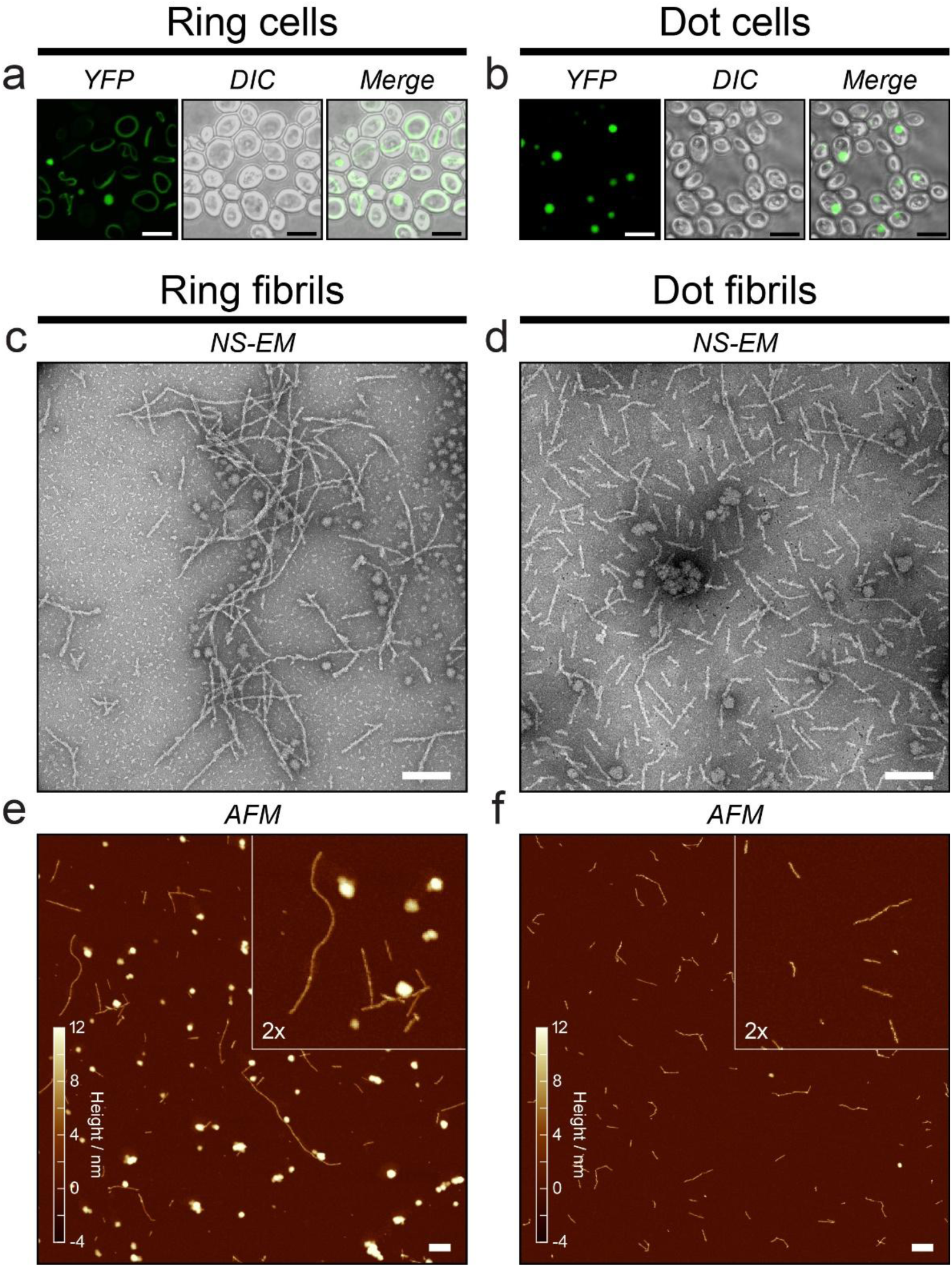
Purification and characterisation of Sup35 Ring and Dot fibrils formed *in vivo*. **a**, Fluorescence microscopy images showing intermediate ring-like aggregates in Ring cells, with a minority of dot-like aggregates also observed. Scale bar, 5µm. **b**, Fluorescence microscopy images showing mature dot-like aggregates in Dot cells. Scale bar, 5µm. **c-f,** Well-dispersed fibrils are observed in the SDS-insoluble extracts purified from Ring and Dot cells. Representative negative-stain EM images of the SDS-resistant fractions purified from Ring (**c**) and Dot (**d**) cells, and representative AFM height-topology images of the corresponding SDS-resistant fractions purified from Ring (**e**) and Dot (**f**) cells, respectively. Scale bar, 200 nm.

Next, insoluble amyloid fibrils were extracted from Ring and Dot cells. Cell lysates were first centrifuged through a 40% sucrose cushion, and the resulting pellets were resuspended in extraction buffer containing 2% SDS before further ultracentrifugation. Immunoblotting confirmed the presence of Sup35 in the extracted SDS-resistant pellets (**Extended Data Fig. 1c**). Semi-denaturing detergent agarose gel electrophoresis (SDD-AGE) further showed that the extracted Ring and Dot Sup35NM aggregates correspond to the highest-molecular-weight SDS-resistant Sup35 species present in the lysates (**Extended Data Fig. 1d**). Direct imaging by negative-stain electron microscopy (NS-EM) (**Fig. 1c-d**) and AFM (**Fig. 1e-f**) revealed dispersed filaments in *ex vivo* material extracted from both Ring and Dot cells. Immunogold labelling with anti-Sup35 and anti-GFP antibodies confirmed that these *ex vivo* filamentous structures are formed by YFP-tagged Sup35NM and further indicate that the C-terminal region of Sup35M containing the anti-Sup35 epitope, as well as the YFP tag, are not incorporated into the ordered fibril core (**Extended Data Fig. 1e**). Together, these results demonstrate that the aggregates observed in Ring and Dot cells contain amyloid fibrils whose ordered cores are formed by Sup35NM (**Fig. 2a**). Hereafter, filaments extracted from Ring cells are referred to as Ring fibrils, and those extracted from Dot cells as Dot fibrils.

**Figure 2.**
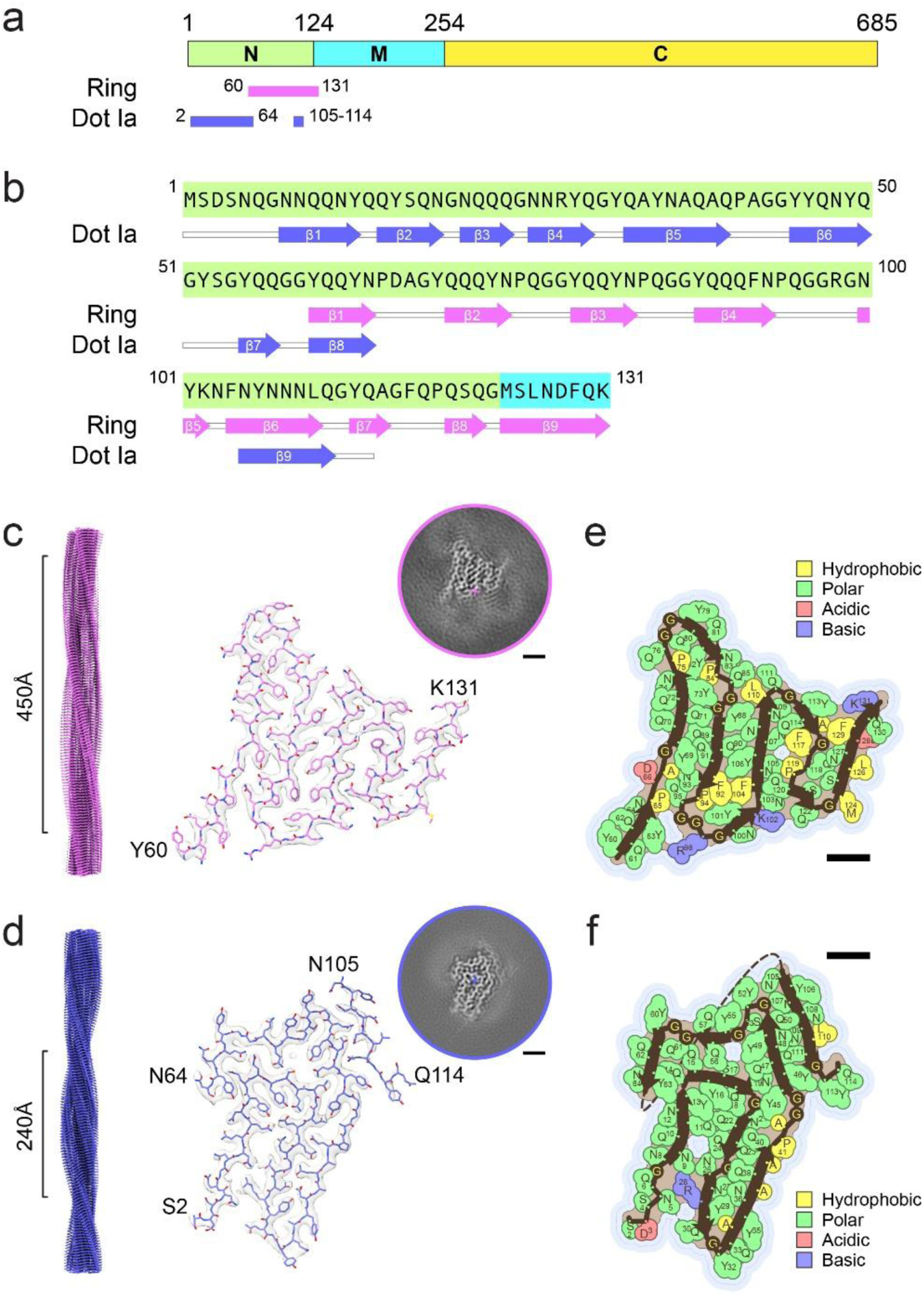
Distinct dominant structures of Sup35 Ring and Dot fibrils resolved by cryo-EM. **a**, Schematic of the Sup35 domain organisation, indicating the ordered core of the dominant Ring fibrils, spanning Y60-K131, and that of the dominant Dot Ia fibrils, spanning S2-N64 and peptide island N105-Q114. **b**, Secondary-structure elements of the amyloid filament cores of *ex vivo* Sup35 Ring and Dot Ia fibrils. Arrows indicate β-strands**. c, d**, Cryo-EM density maps, shown in transparent grey with fitted atomic models, of the Ring fibril (**c**, pink) and Dot Ia fibril (**d**, blue). The 3D reconstructions are also shown in views along the helical axis to the left, with crossover distances indicated. Circular insets show XY cross-sectional views of the 3D reconstructions, with the helical axis indicated by the cross. Scale bar, 10 Å. **e, f**, Cross-sectional schematics of the Ring (**e**) and Dot Ia (**f**) fibril structures, with amino acids coloured according to side-chain chemical functional groups. The images were generated using scripts provided by Michael Sawaya and implemented in the Amyloid Atlas webserver^51^. Scale bar, 10 Å.

### Structural diversity decreases during maturation of polymorphic Sup35 fibril populations

To characterise the *ex vivo* Ring and Dot fibrils, we first analysed the fibril dimensions. Both NS-EM and AFM showed that Ring fibrils are generally longer than Dot fibrils (**Fig. 1c-f**). Quantification of AFM images using Trace_y software^36^ gave a mean length of 150.6 nm for Ring fibrils (n = 634) and 95.6 nm for Dot fibrils (n = 902) (**Extended Data Fig. 2a, b**), consistent with previous *in situ* cryo-tomography showing that Dot fibrils are ∼110 nm long, whereas Ring fibrils can extend to several microns *in vivo*^26^. Most filaments measured 9-13 nm in width in cryo-EM images (**Extended Data Fig. 3a, e**), including the fuzzy coat surrounding the filament core, whereas AFM height measurements, which approximate core width, gave mean values of 5.6 nm for Ring fibrils and 7.0 nm for Dot fibrils (**Extended Data Fig. 2c**). Both were smaller than previously reported widths of Sup35NM-GFP fibrils (∼20-30 nm)^24–26^. This is expected due to the removal of peripheral components such as chaperones during SDS extraction. To assess the biological relevance of these *ex vivo* Sup35NM fibrils, we tested their ability to induce [*PSI^+^*]. Because both particle length and particle number influence transfection efficiency^37^, [*psi⁻*] [*PIN⁺*] yeast cells were transfected with comparable numbers of the *ex vivo* prion fibril particles of similar length, achieved by sonication of Ring fibrils to match the length of Dot fibrils and validated by AFM image analysis (**Extended Data Fig. 2b, c**). Under these conditions, Ring and Dot fibrils induced [*PSI^+^*] with similar efficiency (**Extended Data Fig. 2d, e**). Both Ring and Dot fibrils generated strong and weak [*PSI^+^*] phenotypes, with strong phenotypes predominating, consistent with previous transfection experiments using crude extracts from Ring and Dot cells^24^. These findings confirm that the *ex vivo* fibrils retain biological activity.

Next, we studied the polymorphism in the fibrils by both Cryo-EM (**Fig. 2**, **Extended Data Fig. 3**) and AFM **(Fig. 3)**. Reference-free two-dimensional (2D) classification of cryo-EM images revealed polymorphic fibril populations in both samples, with distinct dominant morphologies in the Ring and Dot states (**Extended Data Fig. 3**). In the Ring sample, 46.8% of particles yielded uninterpretable 2D class averages, probably owing to fibril heterogeneity and poor alignment. 51.7% displayed a twisted morphology with a crossover distance of ∼45 nm, whereas a minor fraction (1.5%) appeared straight, untwisted or had a very long helical pitch (**Extended Data Fig. 3b, c**). By contrast, Dot fibrils showed obvious structural polymorphism with more types of fibrils are identified. Overall, 60.9% of Dot-fibril particles yielded uninterpretable 2D class averages, 31.2% adopted twisted conformations, comprising at least three fibril types (Type I: 25.6%, Type II: 3.1%, and Type III: 2.5%) and 7.9% lacked detectable twist. In total, five distinct fibril types were identified, all different from the dominant Ring fibril type (**Extended Data Fig. 3f, g**). However, all twisted fibrils showed a similar crossover distance of ∼24 nm, suggesting that the structural differences between these polymorphs might be relatively small.

**Figure 3.**
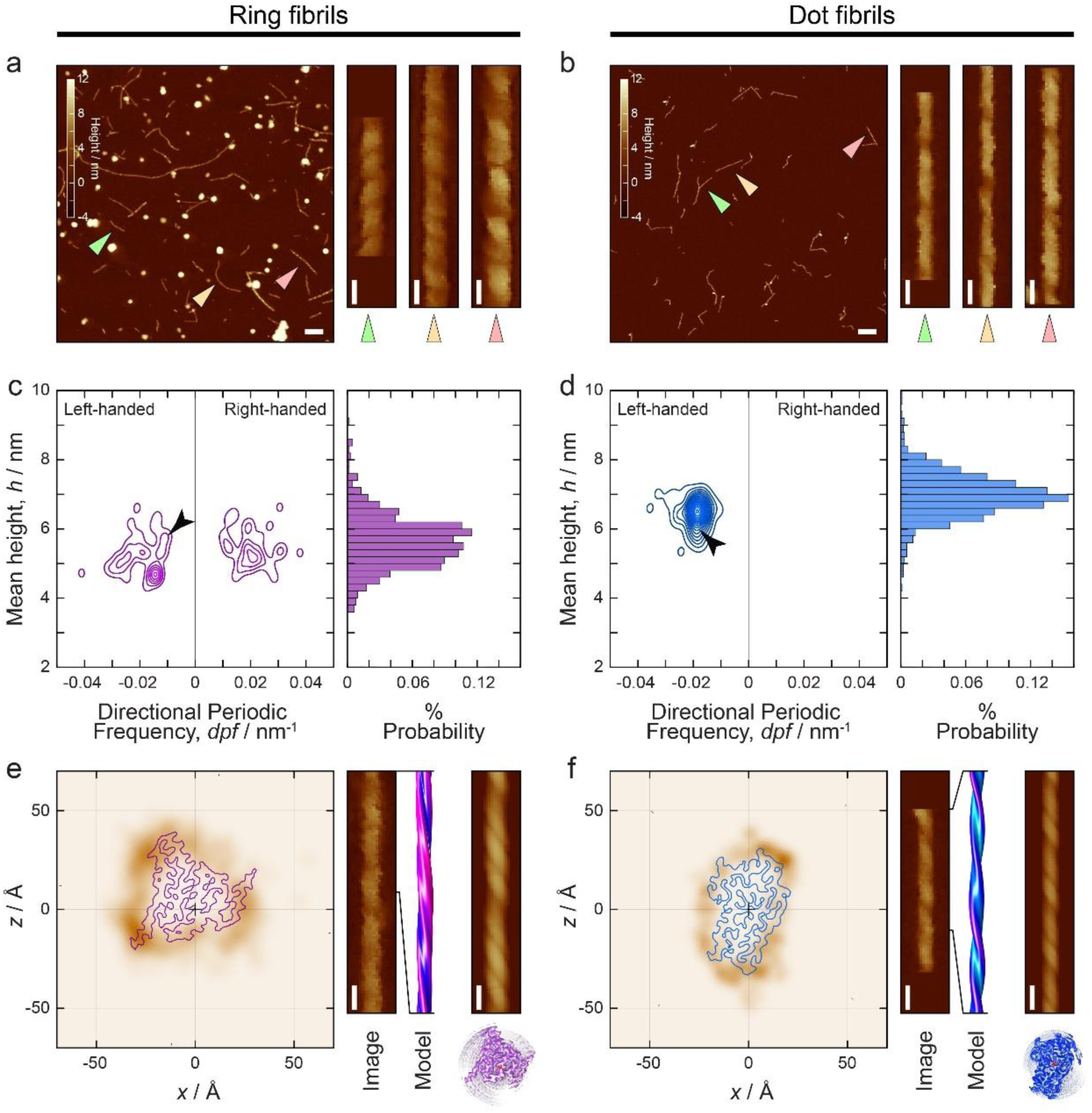
Analysis of structural diversity in Ring and Dot fibril population by CPR-AFM. a, **b**, Representative AFM height-topology images of the ex vivo purified Ring (**a**) and Dot (**b**) fibrils. Scale bar, 200 nm. Digitally straightened examples of individual filaments are shown with 20 nm scalebars, and their positions in the main images are indicated by coloured triangles. **c, d**, Structural diversity of the Ring (**c**) and Dot (**d**) fibril populations visualised as 2D distributions of individual fibril average height, representative of cross-sectional width, and directional periodic frequency (dpf), representative of helical twist. These distributions demonstrate the higher diversity in the Ring fibril population than in the Dot fibril population. The 2D distributions include only fibrils with a clear repeating twist pattern (Ring fibrils, n = 52; Dot fibrils, n = 72), whereas the height distributions to the right include all the well-separated fibrils (Ring fibrils, n = 634; Dot fibrils, n = 902). **e, f**, Cross-sectional CPR-AFM density maps of individual fibrils that best matched the Ring (**e**) and Dot Ia (**f**) structures. The darker brown regions in these cross-sectional density maps indicate areas of contact between the AFM probe tip and the filament cross-section, and are shown in their best-fit orientations together with outlines of the corresponding cross-sections from the cryo-EM density maps. (**d**). Digitally straightened image data, surface-envelope models constructed from the CPR-AFM density maps, and simulated AFM images based on the cryo-EM density maps are also shown for comparison. Scale bar, 20 nm.

Although fibrils from both Ring and Dot states are polymorphic, the extent of structural variation among polymorphs may differ between the two populations. To capture this distinction, we use the term structural diversity to describe the degree of variation among polymorphic structures within a fibril population. To quantify and compare structural diversity at the population level, we analysed individual fibrils by three-dimensional (3D) contact-point reconstruction AFM (CPR-AFM)^36^. This showed that the Ring fibril population is more structurally diverse than the Dot fibril population, as reflected by a broader distribution of polymorphs’ morphometric parameters across all the detected helical filaments (**Fig. 3 a-d, Extended Data Table 2**). Among the twisted fibrils, both left-handed (52%) and right-handed (48%) species were observed in the Ring fibril population, whereas only left-handed twisted fibrils were detected in the Dot fibril population. Overall, structural diversity, quantified as the quadratic mean of all pairwise structural difference scores (d_ξ_)^36^ is 7.96±0.09 for the Ring population, indicating substantially greater diversity than the Dot population, for which the value was 3.31 ± 0.03.

Together, these data provide direct evidence that both the intermediate Ring state and the mature Dot state are polymorphic and comprise clouds of structurally distinct amyloid assemblies. However, the fibril population evolves substantially from the intermediate to the mature state, with structural diversity decreasing during this transition.

### Mature Dot fibrils are structurally distinct from intermediate Ring fibrils

Helical reconstruction of cryo-EM data produced 3D density maps at 3.3 Å and 3.0 Å resolution for the dominant Ring fibril structure (Ring fold) and for Dot Ia, the major conformation within Dot type I fibrils, respectively (**Fig. 2, Extended Data Fig. 3d, h**). Because Ring and Dot fibrils were detected by AFM as predominantly or exclusively left-handed (**Fig. 3c, d**), all the atomic models were built with left-handed helical symmetry. Strikingly, the Ring fold and Dot Ia differ in both conformation and the regions of the Sup35 sequence incorporated into the core. The Ring core comprises residues Y60-K131 and adopts a five-layer meander-shaped fold, whereas Dot Ia comprises residues S2-N64 and N105-Q114 and adopts a four-layer Greek-key fold with an associated peptide island (**Fig. 2c, Extended Data Fig. 4**). Integrative analysis linked both structural maps to the AFM-derived polymorph distribution maps, confirming these structures are represented within the amyloid populations observed by AFM (**Fig. 3e, f**).

Both fibril cores are strongly enriched in hydrophilic residues, with glutamine and asparagine accounting for 41.7% and 47.7% of Ring and Dot Ia folds, respectively (**Fig. 2e, f**). These Q/N-rich segments form extensive hydrogen-bond networks, whereas the less abundant hydrophobic residues are dominated by tyrosine, which contributes hydrogen-bonding, hydrophobic and π-stacking interactions (**Extended Data Fig. 4e-h**). Thus, both folds are stabilised primarily by hydrophilic interactions among neutral polar residues together with tyrosine-mediated hydrophobic stacking (**Extended Data Fig. 4i, j**). Both folds contain nine β-strands, and the two folds partially overlap in sequence at Y60-N64 and N105-Q114 and share conserved β-strand elements within the Y60-N64 and N105-L110 segments (**Fig. 2b**). However, the Ring fold is approximately coplanar, whereas Dot Ia shows substantial height variation within a single layer, spanning 11.5Å along the helical axis from its lowest to highest point (**Extended Data Fig. 4c, d and Fig. 5e**).

In addition to the dominant Dot Ia conformation, cryo-EM 2D and 3D class averages of *ex vivo* Dot fibrils revealed alternative core architectures. 3D classification of type I segments identified an alternative conformation spanning residues Y32-Q40, designated Dot Ib (**Extended Data Fig. 5, 6**). Fewer than 20% of type I segments contributed to Dot Ib classes, and the Dot Ib structure was resolved at 3.1 Å resolution. Relative to Dot Ia, Dot Ib reorganises one extended β-strand into two shorter β-strands through ∼180° rotation of the peptide planes at Y32-Q33 and Q33-A34, shifting the C-terminal region by one helical rise (4.8 Å). We also resolved the structure of type II Dot fibrils (Dot II) at 3.4 Å resolution (**Extended Data Fig. 5, 6**). Dot II adopts the same overall architecture as Dot Ia, but extends eight additional residues at the C terminus and lacks the peptide island spanning N105-Q114 that packs against residues Y46-Y52 in Dot Ia and Ib. In addition, the small density within the hydrophilic cavity seen in Dot Ia and Dot Ib, consistent with an ordered solvent molecule, is absent in Dot II (**Extended Data Fig. 5b, red arrows**). Type III Dot fibrils were also analysed by 3D reconstruction, but no validated 3D density map was obtained. In all Dot folds, R28 is buried within the ordered core, where its positive charge is only partially compensated by hydrogen bonding to backbone peptide groups and the side chain of N9 (**Extended Data Fig. 5f**). Buried positively charged residues have been reported in other amyloid structures, including R293 in TDP-43 filaments from type A FTLD-TDP^38^ and K58 and K80 in *in vitro*-assembled α-synuclein fibrils^39^. In those structures, however, the buried charges are stabilised by post-translational modification or by electrostatic interactions with nearby glutamic acids. This partially compensated buried charge may therefore be energetically unfavourable and may influence stability of the Dot folds. Consistently, thermostability assays showed that Ring fibrils are more resistant to heat-induced disruption than Dot fibrils (**Extended Data Fig. 7**). Together, our high-resolution cryo-EM analysis shows that predominant Dot fibrils adopt 3D architectures distinct from the dominant Ring fibril structure in both conformation and core sequence composition, and contain structural features, including flexible regions outside the ordered core, that may influence their stability and interactions with cellular factors.

### GdnHCl-mediated inhibition of Hsp104 in Dot cells restores the Ring fold and broader structural diversity

Time-lapse microscopy previously showed that, upon GdnHCl treatment, which selectively inhibits ATPase activity of Hsp104^33,34^, caused Dot cells to rapidly and consistently generate daughter cells displaying ring-like aggregates before Ring cells ultimately gave rise to progeny with diffuse fluorescence^24^. To determine whether the same Ring fold structure re-emerges during curing of [*PSI^+^*] from Dot cells, we performed cryo-EM analysis on *ex vivo* fibrils extracted from Dot cells after 20 h of GdnHCl treatment. As expected, GdnHCl induced the curing of [*PSI^+^*], and only 30% of cells still contained aggregates. Among these, the proportion of dot-containing cells decreased, whereas ring-containing cells increased, accounting for 40% and 60% of aggregate-positive cells respectively (**Fig. 4a, b**). The Dot and Ring aggregates were confirmed by Amytracker and ThT staining (**Extended Data Fig. 8a**). The amyloid fibrils from the GdnHCl-treated Dot cells (GTDCs) were extracted and purified using the same method as for Ring and Dot cells (**Fig. 4c, Extended Data Fig. 8b-d**).

**Figure 4.**
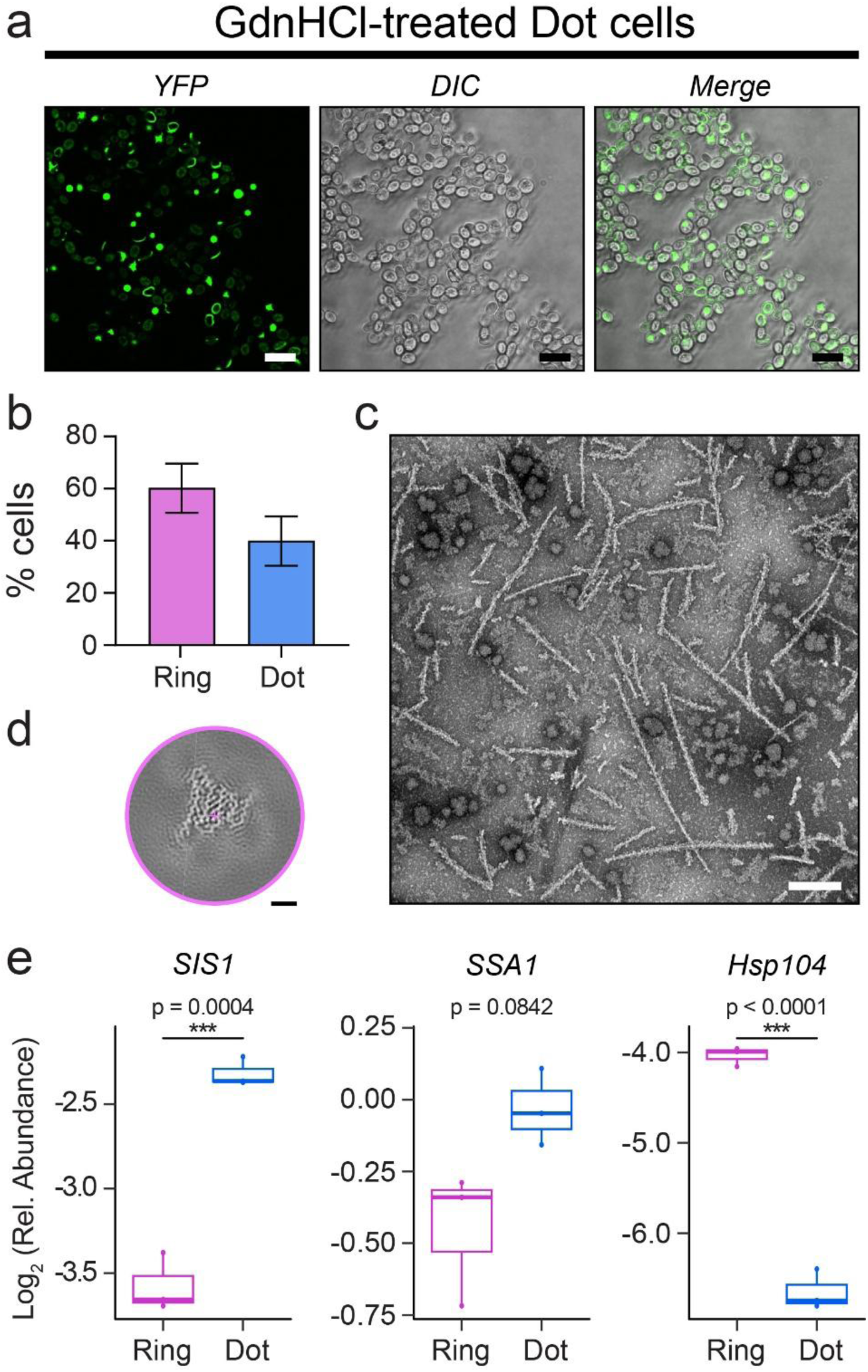
GdnHCl-treated Dot cells restore ring-like aggregates containing Ring-fold fibrils. **a**, Fluorescence microscopy images showing re-emergence of ring-like aggregates in a mature Dot-state population after 20 h of GdnHCl treatment. Scale bar, 10 µm. **b**, Quantification of aggregate morphology in the population shown in **a**. **c**, Negative-stain EM image of fibrils purified from GdnHCl-treated Dot cells (GTDCs). Scale bar, 200 nm. **d**, XY cross-sectional view of the 3D reconstructed cryo-EM map of purified GTDC fibrils. Helical axis is shown as cross. Scale bar, 10 Å. **e**, Quantification of co-purified chaperones by LC–MALDI-TOF/TOF mass spectrometry following anti-GFP pull-down assays from Ring and Dot cell extracts (n=3).

Cryo-EM 2D classification revealed that 51% of all particles yielded uninterpretable 2D class averages, whereas 40% shared the same class averages as twisted Ring filaments, and 9% corresponded to the untwisted Ring fibrils observed in samples extracted from Ring cells. The segments of twisted GTDC fibrils yielded a 3D reconstructed map of 3.4 Å resolution (**Fig. 4d, Extended Data Fig. 8e-i**). Importantly, this map reveals a structure indistinguishable from the Ring fold, as shown by rigid-body fitting of the Ring atomic model into it (**Extended Data Fig. 8f**). Notably, no Dot fibril structures were detected in GTDC fibrils by cryo-EM, although dot-like aggregates were still observed by fluorescence microscopy. These aggregates may therefore contain a small amount of twisted Dot fibrils below the detection limit of cryo-EM, together with increased proportions of irregular or untwisted fibrils, as indicated by 2D classification (**Extended Data Fig. 8e**). GTDC fibril populations were also analysed by AFM (**Extended Data Fig. 9**). Both the length and height distributions of GTDC fibrils closely mirror those of Ring samples (**Extended Data Fig. 9b, d**). This indicates that the GTDC population shows a broader structural diversity, more similar to the Ring fibril population than to the Dot population. Individual-filament AFM analysis provided evidence for both Ring and Dot folds within the GTDC population (**Extended Data Fig. 9e, f**), suggesting that rare twisted Dot fibrils may still be present. In summary, our data show that, as the Dot folds diminish, the Ring fold re-emerges in the fibril population during curing of [*PSI^+^*] by GdnHCl-mediated inhibition of Hsp104, accompanied by restoration of the broader structural diversity characteristic of the Ring state. These findings suggest an important role for Hsp104 in facilitating and maintaining structural evolution of [*PSI^+^*] during maturation.

### Molecular chaperones interact differently to Sup35 Ring and Dot fibrils

Molecular chaperones such as Hsp104, Hsp70 and Hsp40 play key roles in the propagation and maintenance of yeast prions *in vivo*. Chaperones Hsp104, Ssa1 (Hsp70) and Sis1 (Hsp40) have been reported to colocalise with ring- and dot-like aggregates^26^. Because Ring and Dot fibrils incorporate different regions of the Sup35 sequence into their ordered cores, the exposed regions outside the core are likely to differ in their spatial positioning relative to the ordered core, thereby altering accessibility of residues 143-164, the proposed Sis1/Ssa1 binding region^40^. We therefore hypothesised that Sis1 and Ssa1 association would be enhanced with Dot fibrils. To test this, as YFP tag is fused to Sup35 in our yeast cells, we performed anti-GFP pull-down assays from yeast extracts. Co-purifying chaperones associated with Sup35-YFP fibrils were then quantified by LC–MALDI-TOF/TOF mass spectrometry (**Fig. 4e**). Chaperone expression levels were comparable across the Ring and Dot cells (**Extended Data Fig. 8j**). The results show that Dot fibrils captured significantly more Sis1 than Ring fibrils (adjusted p-value = 0.0004). Ssa1 also showed increased association with Dot fibrils, although this did not reach statistical significance after multiple-testing correction (adjusted p-value = 0.0842). In contrast, Hsp104 binding was significantly reduced in Dot fibrils (adjusted p-value < 0.0001), which may reflect more dynamic engagement and thus lower steady-state recovery under these conditions. These results show that Ring and Dot fibrils differentially associate with the chaperone machinery, linking distinct structures to distinct chaperone interactions and further supporting a role for chaperones in structural evolution of [*PSI^+^*].

## Discussion

Structural evolution of Sup35 amyloid fibril populations from the intermediate Ring state to the mature Dot state during the *de novo* formation of [*PSI^+^*] prion *in vivo* is revealed by integrative cryo-EM and AFM analyses. AFM showed that the structural diversity within the amyloid population decreases during maturation. Near-atomic structures show that the Ring amyloid core fold differs from the Dot folds in both conformation and the segments of the Sup35 sequence incorporated into the ordered core. This contrasts with intermediate assembly structures observed in previous time-resolved cryo-EM studies *in vitro* ^11,12^ and *in vivo*^13^, which differ structurally but retain core residue compositions similar to those of their corresponding mature forms. Consistently, cells containing Ring or Dot aggregates give rise only to daughter cells with the corresponding aggregate morphology^24^, in agreement with the structural distinction between the two states. The distinct fibril structures, especially their differences in core sequence composition, are likely to alter filament surface properties and the exposure and spatial positioning of flexible regions outside the core, thereby influencing interactions with cellular factors. Indeed, Ring and Dot fibrils differ in their interactions with chaperones: Dot fibrils show increased binding of the co-chaperones Sis1/Ssa1, together with reduced steady-state recovery of Hsp104. This is consistent with greater exposure of Sis1/Ssa1-binding sites outside the amyloid core in Dot conformers than in the Ring fold. It is also in line with an essential role for Hsp104 in [*PSI^+^*] maturation, as indicated by reversion of the Dot state to the Ring state in both the dominant fibril structure and broader structural diversity of the population during GdnHCl-mediated curing. Together, these findings support a model in which chaperones Sis1, Ssa1 and Hsp104 selectively enrich specific conformers within the Sup35 amyloid population during prion maturation (**Fig. 5**).

**Figure 5.**
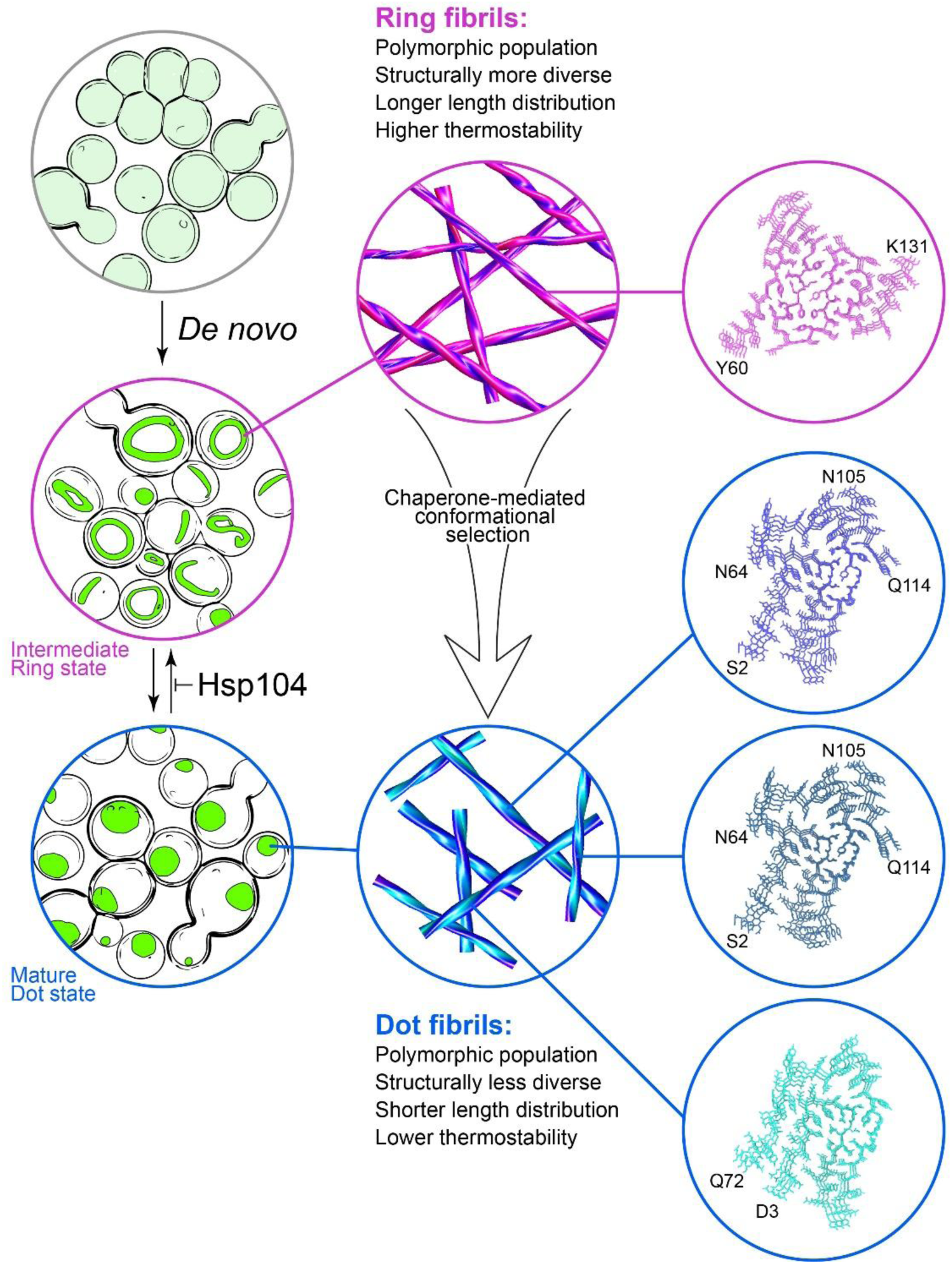
Chaperone-mediated structural evolution during [*PSI^+^*] maturation. Schematic model illustrating the structural evolution of amyloid population during maturation of the [*PSI^+^*] prion from the intermediate Ring to the mature Dot state. Both states contain polymorphic fibril populations, but structural diversity decreases during maturation. Dominant Ring and Dot fibrils differ in both core sequence composition and architecture. They also differ in length, stability and chaperone association. Selective inhibition of Hsp104 by GdnHCl restores the broader structural diversity characteristic of the Ring state and leads to re-emergence of the Ring fold. Together, these data support a model in which, during [*PSI^+^*] prion maturation, molecular chaperones selectively enrich a specific subset of fibril conformers within the Sup35 amyloid population, guiding structural evolution towards conformers that may be particularly permissive to seed-generating chaperone-mediated fragmentation, thereby facilitating propagon production and prion propagation.

Structural comparison showed that both Ring and Dot fibril structures are different from previously reported structures of *in vitro*-formed Sup35 fibrils^41^ and *ex vivo* fibrils formed by a Sup35 variant associated with a strong [*PSI^+^*] phenotype ([*PSI^+^*]-S7)^42^ (**Extended Data Fig. 10**), although all the structures are consistent with the in-register parallel β-sheet architecture of Sup35 fibrils reported before^43^. While no known structure resembles the Dot folds, two *in vitro*-formed Sup35 fibril structures (9XBO, 9XBP) share common substructures with Ring fold at residues 102-131 and 87-129, respectively (**Extended Data Fig. 10c)**, suggesting these regions may contribute to nucleation during *de novo* Sup35 amyloid formation *in vitro* and *in vivo*. Consistently, residues 98-118 have been proposed to form the Sup35 nucleating core^44^. Given that all Ring and Dot amyloid folds, except the minority Dot II polymorph, contain residues 105-114, and a small proportion of dot-like aggregates is detectable at early stages, this suggests that the Dot folds can also arise through self-nucleation, but less efficiently than the Ring fold. Additionally, increasing Sup35 expression preferentially increases the frequency of ring-containing cells relative to dot-containing cells^21^. These observations suggest that the Ring-fold formation is more readily accessed and is therefore likely to be kinetically favoured over the Dot folds during early nucleation. Moreover, the Dot folds contain a buried R28 side chain whose positive charge is only partially compensated, a feature that may contribute to the lower thermal stability of Dot fibrils. The greater average length of Ring fibrils relative to Dot fibrils, both in yeast cells^26^ and after extraction, further suggests that Ring fibrils have higher mechanical stability. Together, these findings raise the possibility that *de novo* formation of the Ring fibril population is favoured over Dot fibrils at early stages of assembly for both kinetic and thermodynamic reasons, consistent with the view that both factors influence the polymorphic distributions observed in *in vitro* aggregation assays^12^. Since multiple factors, including [*PIN^+^*] and Hsp104, may also contribute to heterogeneity during nucleation, they may underlie the greater complexity and structural diversity of the Ring fibril population. The polymorphism in both Ring and Dot Sup35 fibril populations supports the “prion cloud” model, in which prions exist as a dynamic cloud of conformational variants^45,46^, and likely explains the mixed [*PSI^+^*] phenotypes induced by these samples.

During prion maturation, the structural diversity decreases and distinct dominant fibril structures emerge, suggesting a selective pressure for specific conformers during the amyloid evolution *in vivo*. Because site-specific binding of Sis1 and Ssa1 initiates Hsp104-mediated seed-generating fibril fragmentation^40^, the increased association of these co-chaperones to Dot conformers may enhance propagon generation and prion propagation. Dot fibrils are also less thermodynamically and mechanically stable than intermediate Ring fibrils. Previous work showed that increased fibril brittleness enhances prion propagation efficiency^47^, and that GdnHCl treatment inhibits Sup35 fragmentation *in vivo*^48^ and reduces the number of transmissible propagons^33,34^. Together, these findings suggest structural evolution of [*PSI^+^*] *in vivo* towards conformers that may be particularly permissive to seed-generating fragmentation, thereby facilitating propagon production and propagation. This interpretation is consistent with the lower inheritance stability of Ring cells relative to Dot cells, and with the finding that Dot cells, but not Ring cells, can transmit [*PSI^+^*] to mating partners^24^. It is also consistent with our observation that transfection with Ring and Dot fibrils, normalised for particle number and length, yielded similar induction efficiencies, suggesting that intrinsic seeding activity alone does not explain their different behaviours *in vivo* and that additional cellular cofactors are likely to contribute. Although the chaperone interaction patterns and our model are consistent with Hsp104-mediated fragmentation contributing to selection of the Dot fibril conformers, our data do not exclude additional dissolution activities by the chaperone machinery^49^. Importantly, our findings provide structural support for the prion evolutionary theory, in which prion variants are selected based on their ability to propagate or be transmitted^50^.

Both this study and the cryo-EM study of a functional amyloid formed by Orb2 at different time points in *Drosophila*^13^ suggested that cellular components, including molecular chaperones, can shift amyloid dynamics *in vivo*. In contrast to Orb2 amyloid, whose polymorphs become progressively more thermodynamically stable with age, Sup35 amyloid populations shift towards conformers that are less thermodynamically and mechanically stable. This is consistent with our observation that Sup35 amyloid fibrils become shorter and more twisted as they mature from Ring fibrils to Dot fibrils, opposite to the age-dependent progression reported for Orb2 amyloid. This difference may suggest a fundamental distinction between functional and transmissible amyloids: whereas functional amyloids tend to stabilise over time, transmissible amyloid particles such as prions may evolve towards less-stable conformers that favour propagation and transmissibility. An analogous principle of structural evolution may also apply to disease-associated and prion-like amyloids, particularly those formed by proteins containing prion-like domains, such as FUS, TAF15, TDP-43, hnRNPA1 and hnRNPA2/B1, in which specific folds may come to predominate in defined neurodegenerative disorders because they propagate most efficiently under particular cellular conditions, potentially shaped by chaperone-mediated processes.

## Acknowledgements

We thank J. Tyedmers (Heidelberg University) for providing the yeast strains and for helpful suggestions. Cryo-EM data were collected at the ISMB EM facility (Birkbeck College, University of London) with financial support from the Wellcome Trust (202679/Z/16/Z and 206166/Z/17/Z). We are very grateful to N. Lukoyanova and S. Chen for EM support and D. Houldershaw for computing support at Birkbeck. We thank K.Yamashita (University of Tokyo) for help with REFMAC; J. Ribes and M. Patel for help with confocal microscopy; D. Johnson and P. King for infrastructure support; J. Wadsworth, J. Bieschke, P. Kloehn, R. Buckley, A. Wenborn, and M. Batchelor for helpful discussions. We thank R Hervas (University of Hong Kong) for help with SDD-AGE analysis and helpful discussions. This work was funded by the core award to the MRC Prion Unit from the UKRI Medical Research Council (MC_UU_00037/5 to W.Z., studentship to B.A.), and funding from the Biotechnology and Biological Sciences Research Council (BBSRC), UK grant BB/Z516880/1 (SLW and WFX). We acknowledge the University College London Mass Spectrometry Science Technology Platform for mass spectrometry experiments and data analysis.

## Author contributions

Z.W. performed yeast and fibril experiments. Z.W., and B.A. performed chaperone-binding experiments. Z.W., and W.Z. performed cryo-EM and analysed the cryo-EM data. W.Z. and A.G.M. built and refined the atomic models. S.L.W. and W.F.X. carried out AFM experiments and analysed the AFM data. R.Z.C. performed mass spectrometry. R.Z.C. and J.D.G. analysed the data. Z.W., S.L.W., N.K.L., H.R.S., J.C., M.F.T., W.F.X. and W.Z. interpreted the results. W.F.X. and W.Z. supervised the project. All authors contributed to writing the manuscript.

## METHODS

### Yeast Strains, prion induction and maturation

The yeast strain used in this study was the same strain used in previous studies^24,26^. The yeast strain was derived from the 74D-694 background^30^ and carried a deletion of the NM domain in the endogenous *SUP35* gene. A fusion construct encoding Sup35NM (residues 1-253) linked to Yellow Fluorescent Protein (YFP) via an 8-amino-acid linker (GSRVPVEK) at N-termini was stably integrated into the genome under the control of the *GAL1* promoter. To induce *de novo* formation of [*PSI^+^*], yeast cells were inoculated into YPAD medium (1% Yeast Extract, 2% Peptone, 0.1 g L^−1^ Adenine sulfate, 2% D-glucose) at an initial OD₆₀₀ of 0.2 and grown at 30 °C to the early exponential phase (∼4-6 h). Cultures were then centrifuged at 8,000g for 5 min, washed in 1× PBS, and resuspended in YPGal medium (1% Yeast Extract, 2% Peptone, 2% D-galactose, 0.1 g L^−1^ Adenine sulfate) for continued growth for 20 h to induce expression of Sup35NM-YFP by the *GAL1* promoter. This produced mainly ring-like aggregates in yeast cells (Ring cells). To generate mature [*PSI^+^*], the Ring cells were streaked on YPGal plate (1% Yeast Extract, 2% Peptone, 2% D-galactose, 2% Agar) and cultured for 3 days. The cells were then re-streaked three times for ∼100 generations until only dot-like aggregates in the yeast cells (Dot cells) were observed by the fluorescence microscopy. The Dot cells were inoculated into YPGal medium at an initial OD₆₀₀ of 0.2 and grown at 30 °C to mid-log phase (30 h) for fibril extraction. For curing of [*PSI^+^*], Dot cells were treated with guanidine hydrochloride (GdnHCl). Specifically, Dot cells were inoculated into YPGal medium and grown for 6 hours to reach the exponential phase, after which GdnHCl was added to a final concentration of 5 mM, and cells were cultured for an additional 20 hours. The proportions of cells containing ring-like and dot-like aggregates following GdnHCl treatment of Dot cells (GTDCs) were assessed by confocal microscopy.

### Fluorescence Microscopy

Yeast cells were fixed in 4% formaldehyde for 30 min at room temperature. Imaging was performed on a Zeiss LSM 700 confocal microscope using a 488 nm laser for YFP excitation and a 405 nm laser for DAPI excitation, with transmitted light imaging. Images were acquired at 63x magnification with oil immersion optics to obtain high-resolution images of the Sup35NM-YFP aggregates. Z-stack acquisition was performed to capture the three-dimensional organisation of these aggregates, allowing us to distinguish between surface and intracellular localisation. Image processing and analysis were carried out using Zen Blue software.

For thioflavin T (ThT) or Amytracker staining, fixed cells were washed once with PBS and resuspended in 100 µl PBS. Cells were incubated with 15 µM ThT for 20 min at room temperature or with Amytracker 480 (Ebba Biotech AB) at a 1:1,000 dilution overnight at 4 °C. After staining, cells were washed five times with PBS and resuspended in 100 µl PBS. A 2 µl aliquot of stained cells was applied to a high-adhesion microscope slide and air-dried to promote adherence. Slides were sealed with mounting medium and coverslips. ThT and Amytracker 480 fluorescence were imaged using the DAPI channel on a confocal microscope.

### Purification of Sup35 fibrils from yeast cells

The purification procedure was developed from a previously described method^52^. The cells were collected and resuspended in twice their pellet volume of lysis buffer (1× TBS, 10 mM MgCl₂, 0.1% NP-40, one EDTA-free protease inhibitor tablet per 50 ml), slowly dripped into liquid nitrogen for snap-freezing, and stored at −80 °C. Frozen cells were lysed in a pre-chilled Waring blender (Model 8011ESK). The lysate was treated with Benzonase (50 U ml⁻¹) for 15 min at room temperature, followed by the addition of 0.5% NP-40 and centrifugation at 200g for 10 min at 4 °C. The supernatant was overlaid onto a 40% sucrose cushion in lysis buffer and centrifuged at 200,000g for 2 h. Pellets were resuspended in 1 ml lysis buffer supplemented with 2% SDS and centrifuged at 200g for 2 min. The resulting supernatant was centrifuged at 100,000g for 1 h, and pellets were resuspended in 100 µl final buffer (20 mM Tris-HCl, pH 7.5, 100 mM NaCl). Samples were used directly for immunoblotting, immunolabelling and negative-stain EM.

For cryo-EM of fibrils isolated from Ring and Dot cells and transfection assay, 100 µl of suspension was incubated with Benzonase (50 U ml⁻¹) at 37 °C for 15 min with shaking. TBS and 40% sucrose in lysis buffer were added to achieve a final buffer containing 10% sucrose in TBS. The mixture was centrifuged at 100,000g for 1 h at 4 °C, and pellets were resuspended in 100 µl final buffer before centrifugation at 21,000g for 30 min. Supernatants were used for grid preparation. For cryo-EM analysis of GTDC fibrils and all the Ring, Dot and GTDC samples for AFM, the suspension was centrifuged at 21,000g for 15 min at 4 °C and the supernatant was collected. The centrifugation step was then repeated under the same conditions.

### Western blotting and Immunolabelling

Western blotting and Immunogold negative-stain EM were carried out as described^53^. For Western blotting, samples were boiled at 95 °C for 10 min and resolved on Novex 4-20% Tris-glycine gels (Thermo Fisher Scientific). Proteins were transferred to nitrocellulose membranes using a Trans-Blot SD semi-dry transfer cell (Bio-Rad). Primary antibodies included Rabbit anti-Sup35 (MT50, 1:10,000); Rabbit anti-GFP (Thermo Fisher Scientific, A-11122, 1:10,000); Rabbit anti-Hsp104 (Abcam, Ab69549, 1:3,000); Rabbit anti-Sis1 (Cosmo Bio, COP-080051, 1:5,000) and Rabbit anti-alcohol dehydrogenase (Rockland, 200-4143S, 1:5,000). The secondary antibodies included HRP-conjugated Goat anti-rabbit (1:5,000) and IRDye 800CW conjugated Donkey anti-rabbit (LICORbio, 926-32213, 1:5,000).

For immunogold labelling, purified fibril samples were centrifuged at 3,000g for 30 s before grid preparation. A 4 µl aliquot was applied to glow-discharged EM grids, incubated for 40 s, and then blotted with filter paper. Grids were rinsed with 1 ml of gelatine buffer (PBS containing 0.1% gelatine) and blocked by floating on a 50 µl drop of gelatine buffer for 10 min at room temperature. After blotting, grids were incubated for 1 h at room temperature with anti-Sup35 (MT50) or anti-GFP primary antibody at a 1:50 dilution. Following washes with gelatine buffer, grids were incubated for 1 h at room temperature with a secondary anti-rabbit IgG antibody conjugated to 10 nm gold particles (Sigma-Aldrich, G3779, 1:20). Then the grids were negatively stained with NanoW (Nanoprobes) and imaged by TEM.

### Semi-denaturing agarose gel electrophoresis

The Semi-denaturing agarose gel electrophoresis (SDD-AGE) was performed as described previously^48,54^. Samples were incubated in the sample buffer (0.25x TAE, 5% glycerol, 2% SDS and 0.05% bromophenol blue) without boiling and loaded onto a horizontal 1.2% agarose gel prepared in 1x TAE buffer containing 0.1% SDS. The proteins were separated under semi-denaturing conditions and transferred to nitrocellulose membranes by capillary transfer, followed by detection by immunoblotting.

### Yeast transfection with *ex vivo* purified fibrils

Because both particle length and particle number influence transfection efficiency^37^, Ring fibrils were sonicated to match the length distribution of Dot fibrils, which are shorter than 200nm to ensure efficient transfection. Sonication was performed using a probe sonicator (Qsonica Q125) at 20% amplitude with consecutive 5 s on/off cycles on an ice cooled water-bath. Fibril length was monitored by negative-stain EM and AFM. The amount of fibril seeds used for transfection was normalised based on the concentration of Sup35 monomer, determined from Sup35NM-YFP band intensities in Western blots. For the transfection assay, we used the same [*psi^−^*] strain 74D-694 and followed the protocol published before^37,55^. [*psi^−^*] cells were inoculated in 5 ml YEPD and grown overnight at 30 °C, then diluted into 50ml fresh YEPD to an OD600 of 0.2. When the cultures reached an OD600 of 0.6, they were washed and resuspended in 12 ml of ST buffer (1 M sorbitol, 10 mM Tris-HCl pH 7.5). To generate spheroplasts, cells were treated with 600 U of lyticase (Sigma L4025) and 10 mM DTT, and then incubated at 30 °C for 30 min. Spheroplasts were collected by centrifugation (400 g, 5 min), washed with 1.2 M sorbitol and STC buffer (1.2 M Sorbitol, 10 mM Tris-HCl pH 7.5, 10 mM CaCl2), and resuspended in 1 ml STC buffer.

Each transformation reaction consisted of 100 μl spheroplast suspension, 1 μg plasmid DNA (pRS426), 10 μl single-stranded DNA (10 mg ml^−1^) and 10 μl freshly sonicated Sup35NM amyloid fibrils. The mixture was incubated for 10 min at room temperature and gently mixed with 0.9 ml PEG buffer (40% PEG 4000, 10 mM Tris-HCl, pH7.5, 10 mM CaCl_2_). After 30 min of incubation at room temperature, the spheroplasts were collected (400 x g, 5 min) and resuspended in 200 μl SOS medium. Half of the suspension was added to the Top agar (-URA synthetic complete medium containing 2% agar and 1.2 M Sorbitol) and poured onto pre-poured URA-dropout plates. Plates were incubated at 30 °C for 3-4 days. Individual colonies were inoculated into 96-well plates containing YEPD and grown overnight at 30 °C with shaking at 120 rpm. Cultures were replica-plated onto ¼ YEPD (0.25% Yeast Extract, 1% Peptone, 4% Glucose) to assess [*PSI^+^*] phenotypes by colony colour. Ade-dropout plates and ¼ YEPD with 3 mM GdnHCl were used to eliminate any false positives. Three independent transfection assays were performed using different batches of purified Ring and Dot fibrils.

### Thermostability assay

30 μl of Ring or Dot fibril sample was mixed with 60 μl of buffer to give a final reaction mixture containing 20 mM Tris-HCl, pH 7.4, 100 mM NaCl, 1.6% SDS. For each sample, 10 μl aliquots were dispensed into PCR tubes, with eight reactions prepared across a temperature gradient from 25 °C to 95 °C. Samples were incubated for 5 min at the indicated temperature in a PCR thermocycler and then held at 10 °C. Thermal stability was assessed by monitoring the release of soluble Sup35 by western blotting. After incubation, samples were mixed immediately with SDS loading buffer and loaded directly onto SDS-PAGE gels without additional heating. Band intensities were quantified using ImageJ, and data were fitted in GraphPad Prism with a four-parameter logistic model to determine the melting temperature (Tm).

### Immunoprecipitation of Ring and Dot fibrils from yeast cells

The pellet obtained after 40% sucrose-cushion ultracentrifugation of lysates from Ring or Dot cells was resuspended in TBS buffer (20 mM Tris-HCl, pH 7.4, 150 mM NaCl) supplemented with 0.02% NP-40 for immunoprecipitation. Equal amounts of protein were incubated with anti-GFP magnetic beads (ab193983, Abcam) for 1 h at room temperature, washed five times with wash buffer (50 mM Tris-HCl, pH 7.4, 0.5 M NaCl), and then resuspended in 20 µl TBS per reaction for mass spectrometry analysis. Three biological replicates were performed for each fibril type.

### Mass spectrometry and data analysis

To each sample, a lysis buffer (100mM Tris pH 8.5 and 4% SDC) was added; after this step, trypsin was added, and digestion took place overnight. The digestion was stopped, adding 1% trifluoroacetic acid (TFA) to a final pH of 2, SDC was precipitated with centrifugation, and peptides were purified on OASIS HLB plate (Waters). Peptides were dried and dissolved in 0.5% TFA before liquid chromatography-tandem mass spectrometry (MS/MS) analysis. The mixture of tryptic peptides was analysed using a Vanquish NEO high-performance liquid chromatography system coupled online to an Eclipse mass spectrometer (Thermo Fisher Scientific). Buffer A consisted of water acidified with 0.1% formic acid, while buffer B was 80% acetonitrile and 20% water with 0.1% formic acid. The peptides were separated by a 15-cm Waters BEH (1mm internal diameter). The gradient was 3 to 35% B in 52 min at 50 µL/min. Buffer B was then raised to 55% in 2 min and increased to 99% for the cleaning step. Peptides were ionised using a spray voltage of 4 kV and a capillary heated at 320°C. The mass spectrometer was set to acquire full-scan MS spectra (350 to 1400 mass/charge ratio) for a maximum injection time set to Auto at a mass resolution of 120,000 and an automated gain control (AGC) target value of 100%. With a cycle time of 1.2 s, the most intense precursor ions were selected for MS/MS. HCD fragmentation was performed in the HCD cell, with the readout in the Orbitrap mass analyser at a resolution of 30,000 (isolation window of 1.4 Th) and an AGC target value of 200% with a maximum injection time set to Auto and a normalised collision energy of 30%. All raw files were analysed by FragPipe v23 software using the integrated MsFragger v4.3 search engine^56^. All peptides identified were validated with Philosopher v5.1^57^ at 1% FDR and quantified with IonQuant v1.11^58^. All files were searched against the Saccharomyces cerevisiae (strain ATCC 204508 / S288c) Proteome. Msfragger was used with the full-tryptic mode with automatic settings.

For data analysis, proteins assigned to Saccharomyces cerevisiae were retained, and ribosomal proteins were excluded as likely non-specific binders, leaving 228 proteins. As Sup35-YFP was used as the immunoprecipitation bait, protein abundances were normalised to Sup35 abundance to account for differences in bait recovery between samples. Throughout, protein abundance refers to Sup35-normalised abundance.

Protein abundances were processed and group differences were estimated using Auto-Prot version 1.0^59^, implemented with Python v3.11.11 and R v4.4.1. Full details are given at https://github.com/UCL-Biosciences/Auto-Prot. Zero values were treated as missing values, and proteins not detected in at least 66% of samples in all groups were excluded, leaving 160 proteins for downstream analysis. All analyses and visualisations were based on log2-transformed abundances. Differential abundance analysis was carried out using the R package limma (v3.62.2)^60^ including empirical Bayes variance shrinkage and volcano plots were generated using seaborn (v0.12.2)^61^. We used Benjamini-Hochberg False Discovery Rate adjustment to control for multiple testing. Proteins were considered differentially abundant if they showed an FDR-adjusted p-value below 0.05 and an absolute log2 fold change greater than 1.

### AFM specimen preparation, imaging, and analysis

AFM sample specimens were prepared as described previously with modification^37^. Briefly, Ring and Dot samples purified from *S. cerevisiae* cells were diluted 5 or 10 times with sterile-filtered MilliQ water, and 20 µl deposited onto freshly cleaved mica (Agar Scientific) surfaces. Samples were incubated on the discs for 10 minutes before washing with 200 µl of sterile-filtered MilliQ water and dried under a gentle stream of nitrogen gas. A Multimode 8 AFM with a Nanoscope V controller (Bruker) and ScanAsyst probes with nominal tip radius of 2 nm, and nominal spring constant of 0.4 N/m (Bruker) were used for imaging. Images were captured at 1024×1024 px over 4000×4000 or 2048×2048 px over 4000×4000 nm scan size, with structural analysis exclusively performed on the latter (pixel density of 0.51 pixel/nm). Nanoscope Analysis software (version 1.5, Bruker) was used to process all images by baseline flattening the height topology image data to remove tilt and scanner bow. Processed image data were imported into the Trace_y software^62^ for individual filament-level structural analysis, as previously described^63^ to trace, straighten and 3D-reconstruct the fibrils captured in the AFM images. A ‘complete fibrils’ dataset was extracted by tracing along the whole contours of every fibril that were fully visible (e.g. fibrils whose end ran off the image or heavily entangled beyond recognition were not included), regardless of their apparent twist or lack of, for analysis of length and height distribution analysis. A separate dataset containing well-separated fibrils with visible and repeating right- or left-handed helical twist patterns were traced and 3D-reconstructed. Morphometric analysis and integrative structural comparison analysis with cryo-EM maps were carried out according to Aubrey *et al*^36,64^. The quadratic mean of all pairwise structural difference scores (d_ξ_) was calculated using only helical fibrils that yielded 3D reconstructions, because structural comparison of helical and cross-sectional shapes requires reconstructed 3D information. Mean fibril height (nm) and directional periodic frequency were subsequently plotted as 2D contour graphs to visualise the polymorph distributions and differences in the structural diversity of the populations.

### Cryo-electron microscopy

Purified Ring and Dot samples were centrifuged at 3000 g for 30 seconds before being applied on the grids. Cryo-EM grids (Quantifoil 1.2/1.3, 300 mesh) were glow-discharged for 90 seconds. 4µl of Ring or Dot sample was applied to the grids and blotted with filter paper before being plunge-frozen in liquid ethane using a Leica EM GP2 automatic plunge freezer (Leica Microsystems, Germany). The GP2 was set up with the following parameters: sensor blotting, back blotting, additional movement of 0.3mm, blotting time of 6s, humidity of 90% and temperature of 4 ° C.

Cryo-EM data were collected using a 300 kV Titan Krios microscope (Thermo Fisher Scientific) equipped with a K3 direct electron detector operating in super-resolution mode (bin2) with a BioQuantum energy filter (Gatan) operated at a slit width of 10 eV or 20 eV. The data were collected with a nominal magnification of ×105,000 and a pixel size of 0.828 Å. The Ring sample was acquired with 14,002 movies, and the Dot sample was collected with 19,539 movies. Further details are summarised in Extended Data Table 1.

### Helical reconstruction

Datasets were processed in RELION using standard helical reconstruction^9,65^. All the processing was performed in RELION 5.0-beta^66^. Movie frames were gain-corrected, aligned and dose-weighted using RELION’s own implementation of a MontionCor2^67^-like algorithm. Contrast transfer function (CTF) was estimated using CTFFIND4.1^68^. Filaments were picked automatically using Topaz^69,70^. Picked particles were extracted in boxes of 1024 or 512 pixels and downscaled to 256 pixels for initial classification. Reference-free 2D classification, ignoring the CTF until its first peak, was carried out to assess the presence of different types of filaments. Selected particles were re-extracted in boxes of 256 pixels for initial 3D refinement. Initial models were generated *de novo* from 2D class average images using relion_helix_inimodel2d^71^, with an estimated helical rise of 4.75 Å and twist calculated from the crossover distances in the 2D class averages. This initial model was low-pass-filtered to 10 Å before 3D refinement. Subsequent 3D refinement and 3D classification were performed to select particles leading to the best reconstructions, and the helical twist and rise were refined using local searches. Bayesian polishing^72^ and CTF refinement^73^ were applied to increase resolution. Final maps were sharpened using standard post-processing procedures in RELION, and reported resolutions were estimated using a threshold of 0.143 in the Fourier shell correlation (FSC) between two independently refined half-maps^74,75^. We used relion_helix_toolbox to impose helical symmetry on the post-processing maps. Further details are provided in Extended Data Table 1.

### Model building and refinement

Atomic models of the Ring and Dot fibril structures were built de novo in Coot^76^ using the best-resolved maps. Model refinement was performed using Servalcat^77^ and Phenix^78^. In between the rounds of refinement, models were validated using MolProbity^79^, and the clashes and outliers were manually adjusted using ISOLDE^80^ in ChimeraX^81^. For each refined structure, separate model refinements were performed against a single half-map, and the resulting model was compared with the other half-map to confirm the absence of overfitting. Figures were prepared with ChimeraX.

### Reporting summary

Further information on research design is available in the Nature Research Reporting Summary linked to this paper.

## Data availability

Cryo-EM maps have been deposited in the Electron Microscopy Data Bank (EMDB) under accession numbers EMD-57208, EMD-57207, EMD-57209, EMD-57210, EMD-57211 for Ring, Dot Ia, Dot Ib, Dot II, GTDC fibrils, respectively. Corresponding refined atomic models have been deposited in the Protein Data Bank (PDB) under accession numbers 29JH, 29JG, 29JI, 29JJ for Ring, Dot Ia, Dot Ib, Dot II fibrils, respectively. AFM data is shown in the extended data Table. The mass spectrometry proteomics data have been deposited to the ProteomeXchange Consortium via the PRIDE^82^ partner repository with the dataset identifier PXD079885. Any other relevant data is available from the corresponding authors upon reasonable request.

---

**Extended Data Table 1.** Cryo-EM data collection, refinement and validation statistics.

|  | Ring fibrils | Dot fibrils |  |  | GTDC fibrils |
| --- | --- | --- | --- | --- | --- |
| Data collection and processing |  |  |  |  |  |
| Magnification | x105,000 | x105,000 |  |  | x105,000 |
| Voltage (kV) | 300 | 300 |  |  | 300 |
| Electron exposure (e<sup>-</sup>/Å<sup>2</sup>) | 50.16 | 48.22 |  |  | 5.17 |
| Defocus range(μm) | -2.4 to -0.6 | -2.3 to -0.5 |  |  | -2.2 to -0.6 |
| Pixel size (Å) | 0.828 | 0.828 |  |  | 0.828 |
| Micrographs collected | 14,005 | 19,539 |  |  | 20,320 |
| Symmetry imposed | C1 | C1 |  |  | C1 |
| Initial particle images (no.) | 4,143,486 | 1,517,992 |  |  | 2,840,040 |
|  |  | Type Ia | Type Ib | Type II |  |
| Final particle images (no.) | 3,313 | 52,591 | 11,789 | 19,473 | 17,381 |
| Map resolution (Å) | 3.3 | 3.0 | 3.1 | 3.4 | 3.2 |
| Helical twist (°) | -1.915 | -3.24 | -3.51 | -3.32 | -1.936 |
| Helical rise (Å) | 4.81 | 4.72 | 4.72 | 4.71 | 4.81 |
| Refinement |  |  |  |  |  |
| Model resolution(Å) | 3.1 | 3.0 | 3.0 | 3.2 |  |
| FSC threshold | 0.5 | 0.5 | 0.5 | 0.5 |  |
| Map sharpening *B* factor(Å<sup>2</sup>) | -12.99 | -69.70 | -47.70 | -58.51 |  |
| Model composition |  |  |  |  |  |
| Non-hydrogen atoms | 593 | 599 | 599 | 570 |  |
| Protein residues | 72 | 73 | 73 | 70 |  |
| Solvent atoms | 0 | 0 | 0 | 0 |  |
| *B* factor (Å<sup>2</sup>) |  |  |  |  |  |
| Protein | 85.78 | 81.63 | 71.50 | 113.69 |  |
| Solvent | - | - | - | - |  |
| R.m.s. deviations |  |  |  |  |  |
| Bond lengths (Å) | 0.004 | 0.005 | 0.007 | 0.005 |  |
| Bond angles (°) | 0.933 | 0.746 | 1.100 | 1.079 |  |
| Validation |  |  |  |  |  |
| MolProbity score | 1.79 | 1.55 | 1.69 | 1.82 |  |
| Clashscore | 7.26 | 3.67 | 4.59 | 7.72 |  |
| Poor rotamers (%) | 0.00 | 0.00 | 0.00 | 0.00 |  |
| Ramachandran plot |  |  |  |  |  |
| Favored (%) | 94.29 | 94.20 | 92.75 | 94.12 |  |
| Allowed (%) | 5.71 | 5.80 | 7.25 | 5.88 |  |
| Disallowed (%) | 0.0 | 0.00 | 0.00 | 0.00 |  |
| EMDB | 57208 | 57207 | 57209 | 57210 | 57211 |
| PDB | 29JH | 29JG | 29JI | 29JJ |  |

**Extended Data Table 2.**
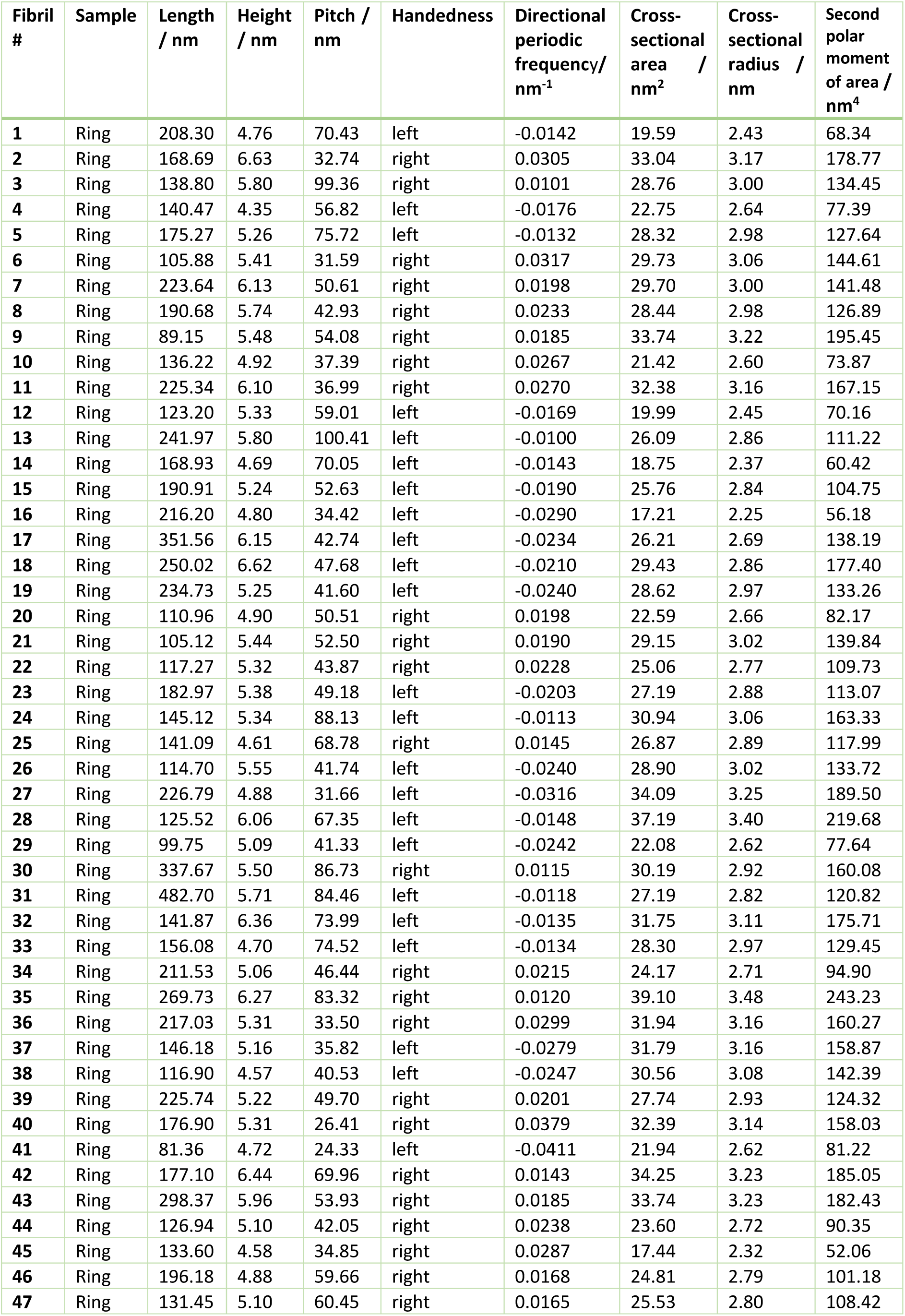
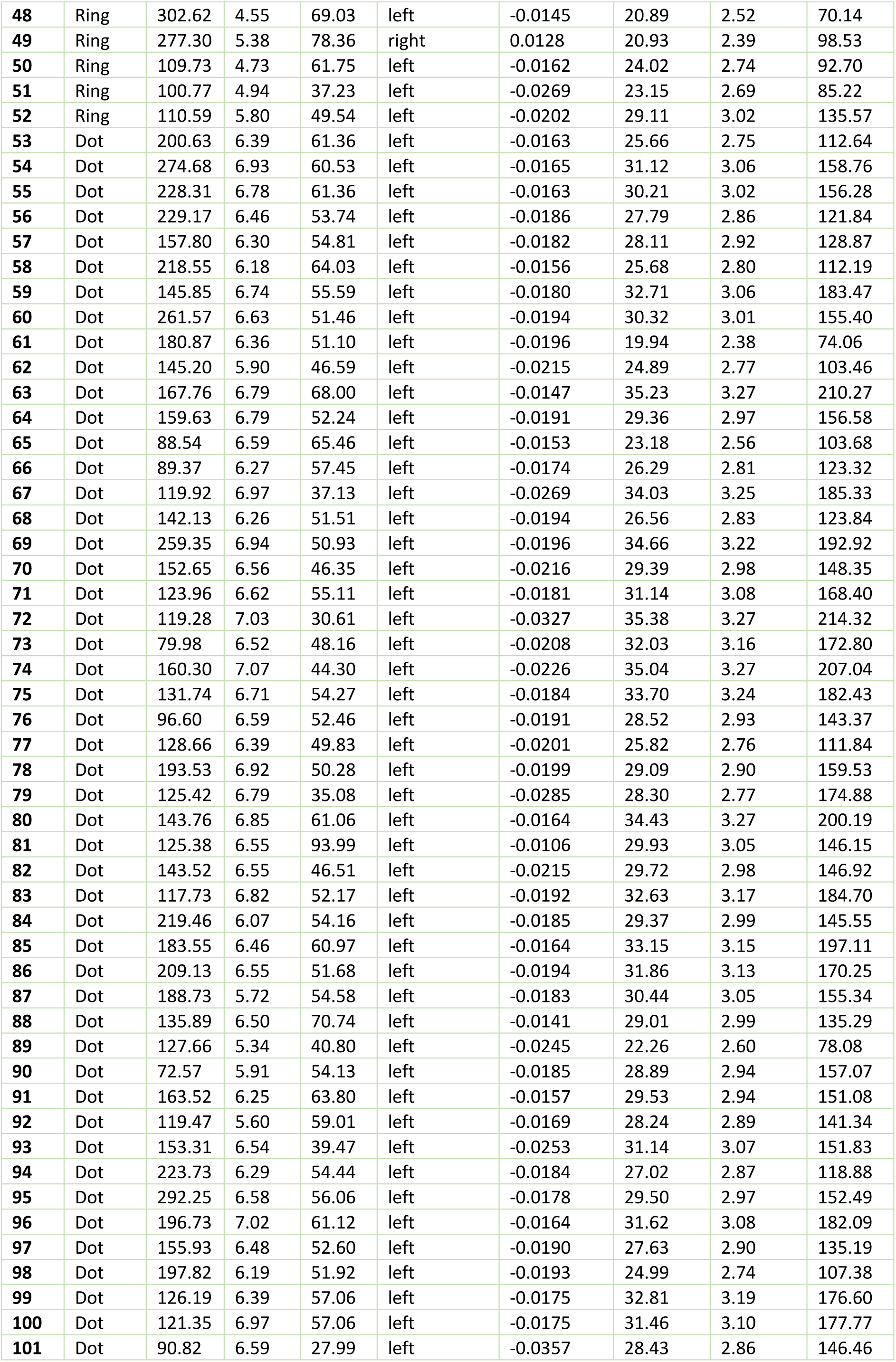
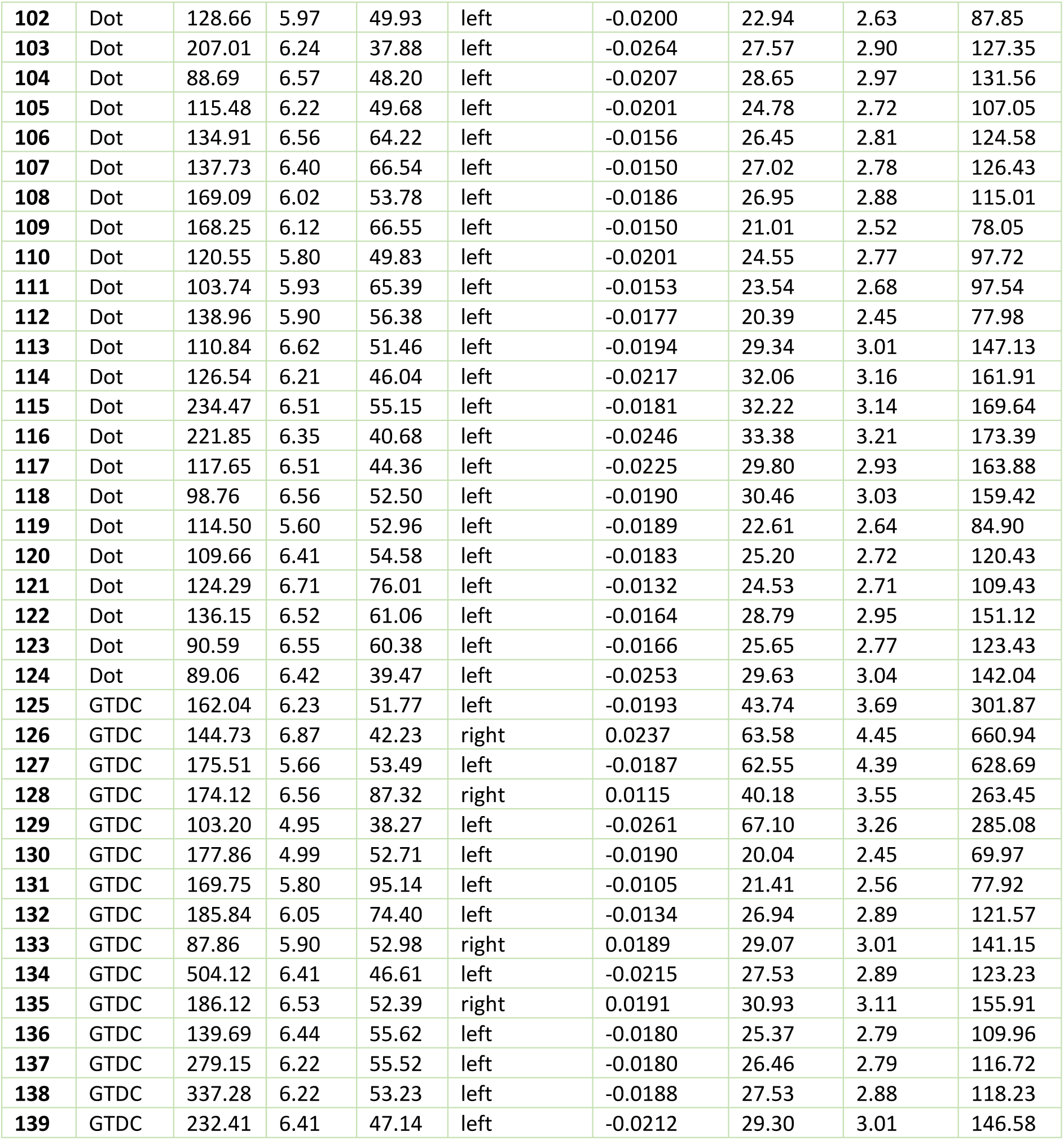
AFM individual filament analysis statistics.

| Fibril # | Sample | Length / nm | Height / nm | Pitch / nm | Handedness | Directional periodic frequency/ nm<sup>-1</sup> | Cross-sectional area / nm<sup>2</sup> | Cross-sectional radius / nm | Second polar moment of area / nm<sup>4</sup> |
| --- | --- | --- | --- | --- | --- | --- | --- | --- | --- |
| 1 | Ring | 208.30 | 4.76 | 70.43 | left | -0.0142 | 19.59 | 2.43 | 68.34 |
| 2 | Ring | 168.69 | 6.63 | 32.74 | right | 0.0305 | 33.04 | 3.17 | 178.77 |
| 3 | Ring | 138.80 | 5.80 | 99.36 | right | 0.0101 | 28.76 | 3.00 | 134.45 |
| 4 | Ring | 140.47 | 4.35 | 56.82 | left | -0.0176 | 22.75 | 2.64 | 77.39 |
| 5 | Ring | 175.27 | 5.26 | 75.72 | left | -0.0132 | 28.32 | 2.98 | 127.64 |
| 6 | Ring | 105.88 | 5.41 | 31.59 | right | 0.0317 | 29.73 | 3.06 | 144.61 |
| 7 | Ring | 223.64 | 6.13 | 50.61 | right | 0.0198 | 29.70 | 3.00 | 141.48 |
| 8 | Ring | 190.68 | 5.74 | 42.93 | right | 0.0233 | 28.44 | 2.98 | 126.89 |
| 9 | Ring | 89.15 | 5.48 | 54.08 | right | 0.0185 | 33.74 | 3.22 | 195.45 |
| 10 | Ring | 136.22 | 4.92 | 37.39 | right | 0.0267 | 21.42 | 2.60 | 73.87 |
| 11 | Ring | 225.34 | 6.10 | 36.99 | right | 0.0270 | 32.38 | 3.16 | 167.15 |
| 12 | Ring | 123.20 | 5.33 | 59.01 | left | -0.0169 | 19.99 | 2.45 | 70.16 |
| 13 | Ring | 241.97 | 5.80 | 100.41 | left | -0.0100 | 26.09 | 2.86 | 111.22 |
| 14 | Ring | 168.93 | 4.69 | 70.05 | left | -0.0143 | 18.75 | 2.37 | 60.42 |
| 15 | Ring | 190.91 | 5.24 | 52.63 | left | -0.0190 | 25.76 | 2.84 | 104.75 |
| 16 | Ring | 216.20 | 4.80 | 34.42 | left | -0.0290 | 17.21 | 2.25 | 56.18 |
| 17 | Ring | 351.56 | 6.15 | 42.74 | left | -0.0234 | 26.21 | 2.69 | 138.19 |
| 18 | Ring | 250.02 | 6.62 | 47.68 | left | -0.0210 | 29.43 | 2.86 | 177.40 |
| 19 | Ring | 234.73 | 5.25 | 41.60 | left | -0.0240 | 28.62 | 2.97 | 133.26 |
| 20 | Ring | 110.96 | 4.90 | 50.51 | right | 0.0198 | 22.59 | 2.66 | 82.17 |
| 21 | Ring | 105.12 | 5.44 | 52.50 | right | 0.0190 | 29.15 | 3.02 | 139.84 |
| 22 | Ring | 117.27 | 5.32 | 43.87 | right | 0.0228 | 25.06 | 2.77 | 109.73 |
| 23 | Ring | 182.97 | 5.38 | 49.18 | left | -0.0203 | 27.19 | 2.88 | 113.07 |
| 24 | Ring | 145.12 | 5.34 | 88.13 | left | -0.0113 | 30.94 | 3.06 | 163.33 |
| 25 | Ring | 141.09 | 4.61 | 68.78 | right | 0.0145 | 26.87 | 2.89 | 117.99 |
| 26 | Ring | 114.70 | 5.55 | 41.74 | left | -0.0240 | 28.90 | 3.02 | 133.72 |
| 27 | Ring | 226.79 | 4.88 | 31.66 | left | -0.0316 | 34.09 | 3.25 | 189.50 |
| 28 | Ring | 125.52 | 6.06 | 67.35 | left | -0.0148 | 37.19 | 3.40 | 219.68 |
| 29 | Ring | 99.75 | 5.09 | 41.33 | left | -0.0242 | 22.08 | 2.62 | 77.64 |
| 30 | Ring | 337.67 | 5.50 | 86.73 | right | 0.0115 | 30.19 | 2.92 | 160.08 |
| 31 | Ring | 482.70 | 5.71 | 84.46 | left | -0.0118 | 27.19 | 2.82 | 120.82 |
| 32 | Ring | 141.87 | 6.36 | 73.99 | left | -0.0135 | 31.75 | 3.11 | 175.71 |
| 33 | Ring | 156.08 | 4.70 | 74.52 | left | -0.0134 | 28.30 | 2.97 | 129.45 |
| 34 | Ring | 211.53 | 5.06 | 46.44 | right | 0.0215 | 24.17 | 2.71 | 94.90 |
| 35 | Ring | 269.73 | 6.27 | 83.32 | right | 0.0120 | 39.10 | 3.48 | 243.23 |
| 36 | Ring | 217.03 | 5.31 | 33.50 | right | 0.0299 | 31.94 | 3.16 | 160.27 |
| 37 | Ring | 146.18 | 5.16 | 35.82 | left | -0.0279 | 31.79 | 3.16 | 158.87 |
| 38 | Ring | 116.90 | 4.57 | 40.53 | left | -0.0247 | 30.56 | 3.08 | 142.39 |
| 39 | Ring | 225.74 | 5.22 | 49.70 | right | 0.0201 | 27.74 | 2.93 | 124.32 |
| 40 | Ring | 176.90 | 5.31 | 26.41 | right | 0.0379 | 32.39 | 3.14 | 158.03 |
| 41 | Ring | 81.36 | 4.72 | 24.33 | left | -0.0411 | 21.94 | 2.62 | 81.22 |
| 42 | Ring | 177.10 | 6.44 | 69.96 | right | 0.0143 | 34.25 | 3.23 | 185.05 |
| 43 | Ring | 298.37 | 5.96 | 53.93 | right | 0.0185 | 33.74 | 3.23 | 182.43 |
| 44 | Ring | 126.94 | 5.10 | 42.05 | right | 0.0238 | 23.60 | 2.72 | 90.35 |
| 45 | Ring | 133.60 | 4.58 | 34.85 | right | 0.0287 | 17.44 | 2.32 | 52.06 |
| 46 | Ring | 196.18 | 4.88 | 59.66 | right | 0.0168 | 24.81 | 2.79 | 101.18 |
| 47 | Ring | 131.45 | 5.10 | 60.45 | right | 0.0165 | 25.53 | 2.80 | 108.42 |
| 48 | Ring | 302.62 | 4.55 | 69.03 | left | -0.0145 | 20.89 | 2.52 | 70.14 |
| 49 | Ring | 277.30 | 5.38 | 78.36 | right | 0.0128 | 20.93 | 2.39 | 98.53 |
| 50 | Ring | 109.73 | 4.73 | 61.75 | left | -0.0162 | 24.02 | 2.74 | 92.70 |
| 51 | Ring | 100.77 | 4.94 | 37.23 | left | -0.0269 | 23.15 | 2.69 | 85.22 |
| 52 | Ring | 110.59 | 5.80 | 49.54 | left | -0.0202 | 29.11 | 3.02 | 135.57 |
| 53 | Dot | 200.63 | 6.39 | 61.36 | left | -0.0163 | 25.66 | 2.75 | 112.64 |
| 54 | Dot | 274.68 | 6.93 | 60.53 | left | -0.0165 | 31.12 | 3.06 | 158.76 |
| 55 | Dot | 228.31 | 6.78 | 61.36 | left | -0.0163 | 30.21 | 3.02 | 156.28 |
| 56 | Dot | 229.17 | 6.46 | 53.74 | left | -0.0186 | 27.79 | 2.86 | 121.84 |
| 57 | Dot | 157.80 | 6.30 | 54.81 | left | -0.0182 | 28.11 | 2.92 | 128.87 |
| 58 | Dot | 218.55 | 6.18 | 64.03 | left | -0.0156 | 25.68 | 2.80 | 112.19 |
| 59 | Dot | 145.85 | 6.74 | 55.59 | left | -0.0180 | 32.71 | 3.06 | 183.47 |
| 60 | Dot | 261.57 | 6.63 | 51.46 | left | -0.0194 | 30.32 | 3.01 | 155.40 |
| 61 | Dot | 180.87 | 6.36 | 51.10 | left | -0.0196 | 19.94 | 2.38 | 74.06 |
| 62 | Dot | 145.20 | 5.90 | 46.59 | left | -0.0215 | 24.89 | 2.77 | 103.46 |
| 63 | Dot | 167.76 | 6.79 | 68.00 | left | -0.0147 | 35.23 | 3.27 | 210.27 |
| 64 | Dot | 159.63 | 6.79 | 52.24 | left | -0.0191 | 29.36 | 2.97 | 156.58 |
| 65 | Dot | 88.54 | 6.59 | 65.46 | left | -0.0153 | 23.18 | 2.56 | 103.68 |
| 66 | Dot | 89.37 | 6.27 | 57.45 | left | -0.0174 | 26.29 | 2.81 | 123.32 |
| 67 | Dot | 119.92 | 6.97 | 37.13 | left | -0.0269 | 34.03 | 3.25 | 185.33 |
| 68 | Dot | 142.13 | 6.26 | 51.51 | left | -0.0194 | 26.56 | 2.83 | 123.84 |
| 69 | Dot | 259.35 | 6.94 | 50.93 | left | -0.0196 | 34.66 | 3.22 | 192.92 |
| 70 | Dot | 152.65 | 6.56 | 46.35 | left | -0.0216 | 29.39 | 2.98 | 148.35 |
| 71 | Dot | 123.96 | 6.62 | 55.11 | left | -0.0181 | 31.14 | 3.08 | 168.40 |
| 72 | Dot | 119.28 | 7.03 | 30.61 | left | -0.0327 | 35.38 | 3.27 | 214.32 |
| 73 | Dot | 79.98 | 6.52 | 48.16 | left | -0.0208 | 32.03 | 3.16 | 172.80 |
| 74 | Dot | 160.30 | 7.07 | 44.30 | left | -0.0226 | 35.04 | 3.27 | 207.04 |
| 75 | Dot | 131.74 | 6.71 | 54.27 | left | -0.0184 | 33.70 | 3.24 | 182.43 |
| 76 | Dot | 96.60 | 6.59 | 52.46 | left | -0.0191 | 28.52 | 2.93 | 143.37 |
| 77 | Dot | 128.66 | 6.39 | 49.83 | left | -0.0201 | 25.82 | 2.76 | 111.84 |
| 78 | Dot | 193.53 | 6.92 | 50.28 | left | -0.0199 | 29.09 | 2.90 | 159.53 |
| 79 | Dot | 125.42 | 6.79 | 35.08 | left | -0.0285 | 28.30 | 2.77 | 174.88 |
| 80 | Dot | 143.76 | 6.85 | 61.06 | left | -0.0164 | 34.43 | 3.27 | 200.19 |
| 81 | Dot | 125.38 | 6.55 | 93.99 | left | -0.0106 | 29.93 | 3.05 | 146.15 |
| 82 | Dot | 143.52 | 6.55 | 46.51 | left | -0.0215 | 29.72 | 2.98 | 146.92 |
| 83 | Dot | 117.73 | 6.82 | 52.17 | left | -0.0192 | 32.63 | 3.17 | 184.70 |
| 84 | Dot | 219.46 | 6.07 | 54.16 | left | -0.0185 | 29.37 | 2.99 | 145.55 |
| 85 | Dot | 183.55 | 6.46 | 60.97 | left | -0.0164 | 33.15 | 3.15 | 197.11 |
| 86 | Dot | 209.13 | 6.55 | 51.68 | left | -0.0194 | 31.86 | 3.13 | 170.25 |
| 87 | Dot | 188.73 | 5.72 | 54.58 | left | -0.0183 | 30.44 | 3.05 | 155.34 |
| 88 | Dot | 135.89 | 6.50 | 70.74 | left | -0.0141 | 29.01 | 2.99 | 135.29 |
| 89 | Dot | 127.66 | 5.34 | 40.80 | left | -0.0245 | 22.26 | 2.60 | 78.08 |
| 90 | Dot | 72.57 | 5.91 | 54.13 | left | -0.0185 | 28.89 | 2.94 | 157.07 |
| 91 | Dot | 163.52 | 6.25 | 63.80 | left | -0.0157 | 29.53 | 2.94 | 151.08 |
| 92 | Dot | 119.47 | 5.60 | 59.01 | left | -0.0169 | 28.24 | 2.89 | 141.34 |
| 93 | Dot | 153.31 | 6.54 | 39.47 | left | -0.0253 | 31.14 | 3.07 | 151.83 |
| 94 | Dot | 223.73 | 6.29 | 54.44 | left | -0.0184 | 27.02 | 2.87 | 118.88 |
| 95 | Dot | 292.25 | 6.58 | 56.06 | left | -0.0178 | 29.50 | 2.97 | 152.49 |
| 96 | Dot | 196.73 | 7.02 | 61.12 | left | -0.0164 | 31.62 | 3.08 | 182.09 |
| 97 | Dot | 155.93 | 6.48 | 52.60 | left | -0.0190 | 27.63 | 2.90 | 135.19 |
| 98 | Dot | 197.82 | 6.19 | 51.92 | left | -0.0193 | 24.99 | 2.74 | 107.38 |
| 99 | Dot | 126.19 | 6.39 | 57.06 | left | -0.0175 | 32.81 | 3.19 | 176.60 |
| 100 | Dot | 121.35 | 6.97 | 57.06 | left | -0.0175 | 31.46 | 3.10 | 177.77 |
| 101 | Dot | 90.82 | 6.59 | 27.99 | left | -0.0357 | 28.43 | 2.86 | 146.46 |
| 102 | Dot | 128.66 | 5.97 | 49.93 | left | -0.0200 | 22.94 | 2.63 | 87.85 |
| 103 | Dot | 207.01 | 6.24 | 37.88 | left | -0.0264 | 27.57 | 2.90 | 127.35 |
| 104 | Dot | 88.69 | 6.57 | 48.20 | left | -0.0207 | 28.65 | 2.97 | 131.56 |
| 105 | Dot | 115.48 | 6.22 | 49.68 | left | -0.0201 | 24.78 | 2.72 | 107.05 |
| 106 | Dot | 134.91 | 6.56 | 64.22 | left | -0.0156 | 26.45 | 2.81 | 124.58 |
| 107 | Dot | 137.73 | 6.40 | 66.54 | left | -0.0150 | 27.02 | 2.78 | 126.43 |
| 108 | Dot | 169.09 | 6.02 | 53.78 | left | -0.0186 | 26.95 | 2.88 | 115.01 |
| 109 | Dot | 168.25 | 6.12 | 66.55 | left | -0.0150 | 21.01 | 2.52 | 78.05 |
| 110 | Dot | 120.55 | 5.80 | 49.83 | left | -0.0201 | 24.55 | 2.77 | 97.72 |
| 111 | Dot | 103.74 | 5.93 | 65.39 | left | -0.0153 | 23.54 | 2.68 | 97.54 |
| 112 | Dot | 138.96 | 5.90 | 56.38 | left | -0.0177 | 20.39 | 2.45 | 77.98 |
| 113 | Dot | 110.84 | 6.62 | 51.46 | left | -0.0194 | 29.34 | 3.01 | 147.13 |
| 114 | Dot | 126.54 | 6.21 | 46.04 | left | -0.0217 | 32.06 | 3.16 | 161.91 |
| 115 | Dot | 234.47 | 6.51 | 55.15 | left | -0.0181 | 32.22 | 3.14 | 169.64 |
| 116 | Dot | 221.85 | 6.35 | 40.68 | left | -0.0246 | 33.38 | 3.21 | 173.39 |
| 117 | Dot | 117.65 | 6.51 | 44.36 | left | -0.0225 | 29.80 | 2.93 | 163.88 |
| 118 | Dot | 98.76 | 6.56 | 52.50 | left | -0.0190 | 30.46 | 3.03 | 159.42 |
| 119 | Dot | 114.50 | 5.60 | 52.96 | left | -0.0189 | 22.61 | 2.64 | 84.90 |
| 120 | Dot | 109.66 | 6.41 | 54.58 | left | -0.0183 | 25.20 | 2.72 | 120.43 |
| 121 | Dot | 124.29 | 6.71 | 76.01 | left | -0.0132 | 24.53 | 2.71 | 109.43 |
| 122 | Dot | 136.15 | 6.52 | 61.06 | left | -0.0164 | 28.79 | 2.95 | 151.12 |
| 123 | Dot | 90.59 | 6.55 | 60.38 | left | -0.0166 | 25.65 | 2.77 | 123.43 |
| 124 | Dot | 89.06 | 6.42 | 39.47 | left | -0.0253 | 29.63 | 3.04 | 142.04 |
| 125 | GTDC | 162.04 | 6.23 | 51.77 | left | -0.0193 | 43.74 | 3.69 | 301.87 |
| 126 | GTDC | 144.73 | 6.87 | 42.23 | right | 0.0237 | 63.58 | 4.45 | 660.94 |
| 127 | GTDC | 175.51 | 5.66 | 53.49 | left | -0.0187 | 62.55 | 4.39 | 628.69 |
| 128 | GTDC | 174.12 | 6.56 | 87.32 | right | 0.0115 | 40.18 | 3.55 | 263.45 |
| 129 | GTDC | 103.20 | 4.95 | 38.27 | left | -0.0261 | 67.10 | 3.26 | 285.08 |
| 130 | GTDC | 177.86 | 4.99 | 52.71 | left | -0.0190 | 20.04 | 2.45 | 69.97 |
| 131 | GTDC | 169.75 | 5.80 | 95.14 | left | -0.0105 | 21.41 | 2.56 | 77.92 |
| 132 | GTDC | 185.84 | 6.05 | 74.40 | left | -0.0134 | 26.94 | 2.89 | 121.57 |
| 133 | GTDC | 87.86 | 5.90 | 52.98 | right | 0.0189 | 29.07 | 3.01 | 141.15 |
| 134 | GTDC | 504.12 | 6.41 | 46.61 | left | -0.0215 | 27.53 | 2.89 | 123.23 |
| 135 | GTDC | 186.12 | 6.53 | 52.39 | right | 0.0191 | 30.93 | 3.11 | 155.91 |
| 136 | GTDC | 139.69 | 6.44 | 55.62 | left | -0.0180 | 25.37 | 2.79 | 109.96 |
| 137 | GTDC | 279.15 | 6.22 | 55.52 | left | -0.0180 | 26.46 | 2.79 | 116.72 |
| 138 | GTDC | 337.28 | 6.22 | 53.23 | left | -0.0188 | 27.53 | 2.88 | 118.23 |
| 139 | GTDC | 232.41 | 6.41 | 47.14 | left | -0.0212 | 29.30 | 3.01 | 146.58 |

## EXTENDED DATA FIGURES

**Extended Data Fig. 1.**
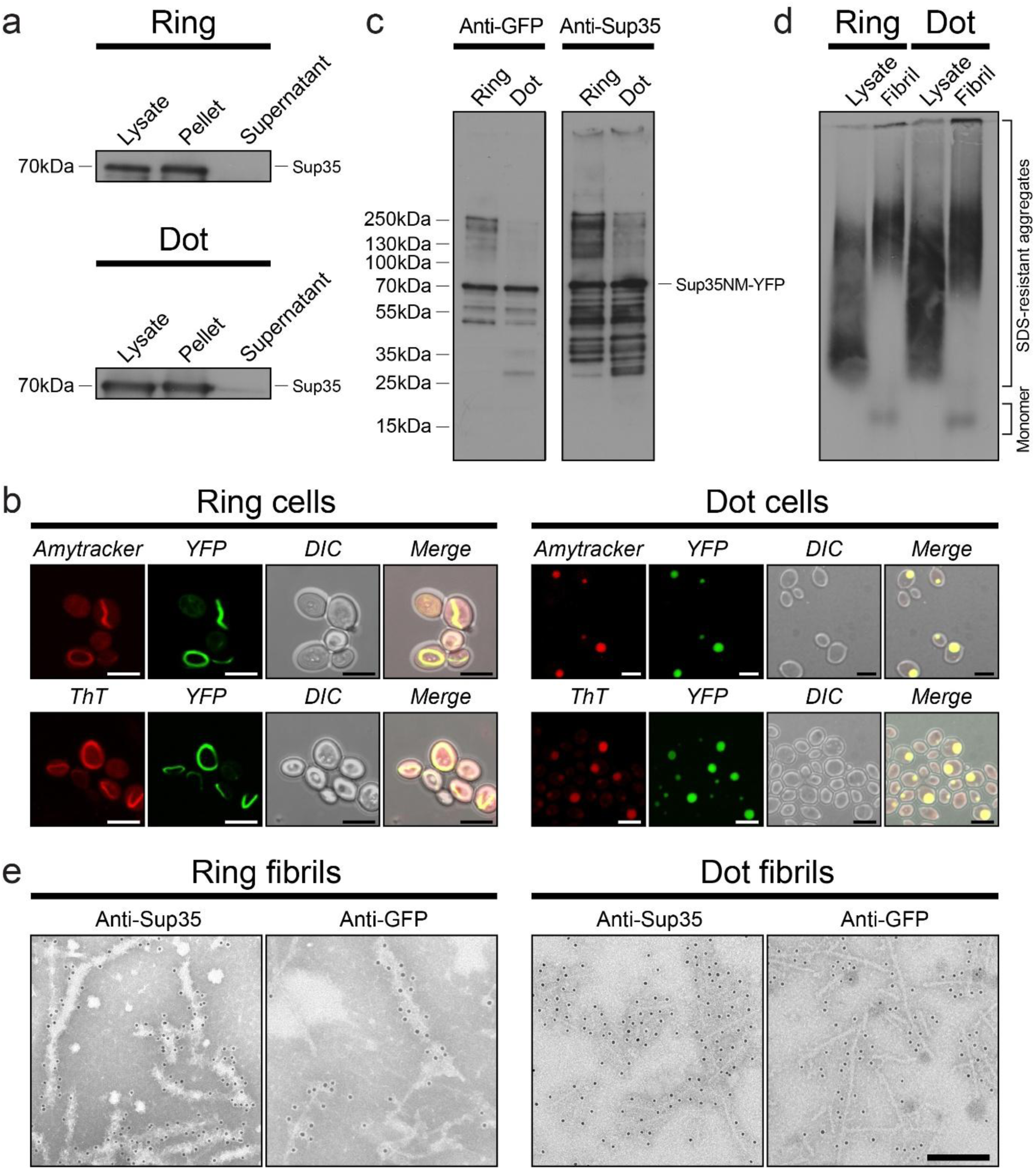
Further characterisation of Sup35 Ring and Dot fibrils. **a**, Sedimentation assays of lysates from Ring and Dot cells showing Sup35 predominantly formed aggregates. Antibody: anti-Sup35. **b**, Fluorescence microscopy images showing Amytracker and ThT colocalised with ring-like and dot-like aggregates in both Ring and Dot cells. Scale bar, 5 µm. **c**, Immunoblots of extracted Ring and Dot fibrils with antibodies anti-GFP and anti-Sup35. **d**, SDD-AGE of Ring and Dot fibrils followed by immunoblotting with anti-Sup35 antibody. **e**, Immunogold negative-stain EM images of Ring and Dot fibrils with antibodies anti-Sup35 and anti-GFP. Scale bar, 200 nm.

**Extended Data Fig. 2.**
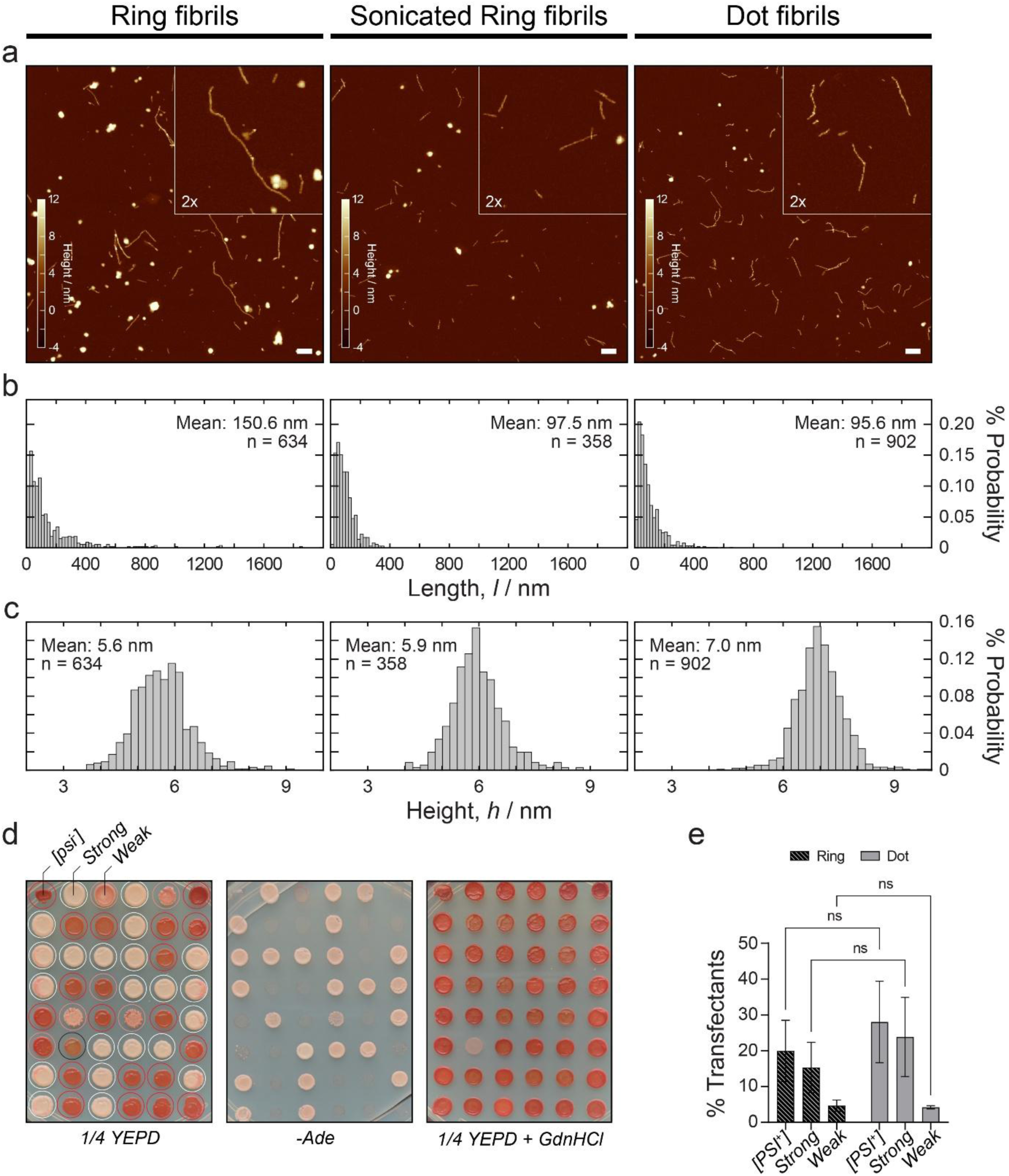
Infectivity assay of Sup35 Ring and Dot fibrils. **a**, Representative AFM height-topology images of Sup35 Ring fibrils, sonicated Ring fibrils and Dot fibrils. **b**, Length distribution of Sup35 Ring fibrils, sonicated Ring fibrils and Dot fibrils. **c**, Height-distribution of Sup35 Ring fibrils, sonicated Ring fibrils and Dot fibrils. **d**, Representative images illustrating the efficiency of Dot fibrils in converting [*psi^−^*] yeast cells to [*PSI^+^*] cells upon transfection. Negative [*psi^−^*] and positive [*PSI^+^*] strong and weak phenotypes are indicated with red, white and pink circles, respectively. The colony which didn’t turn red in the plate of 1/4YEPD supplemented with 3 mM GdnHCl is indicated with a black circle and excluded from the statistics. **e,** Quantification and comparison of the transfection efficiency of Sup35 Ring and Dot fibrils (n=3).

**Extended Data Fig. 3.**
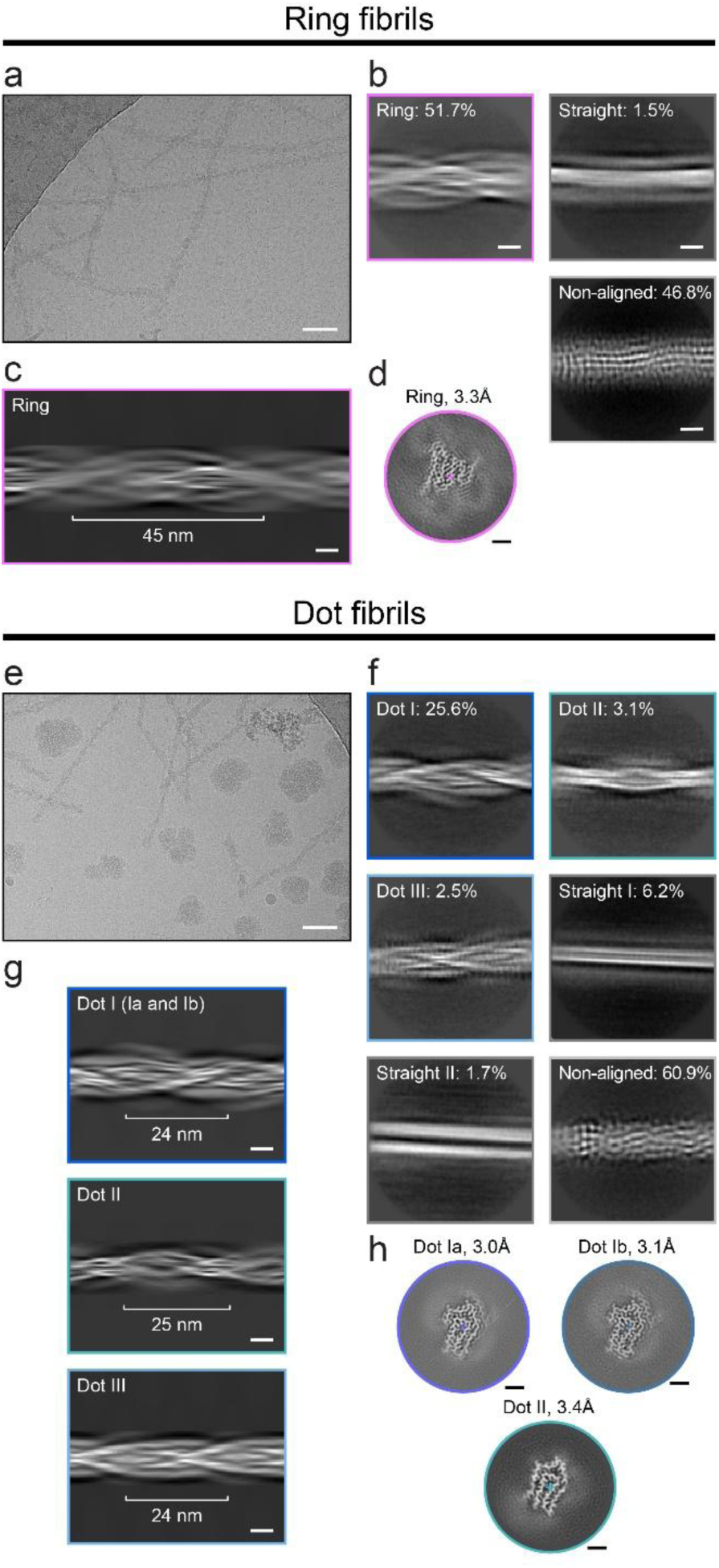
Cryo-EM study of Sup35 Ring and Dot fibrils. **a**, Representative cryo-EM micrograph of Ring fibrils. Scale bar, 50 nm. **b**, Representative reference-free 2D class averages of segments from the cryo-EM dataset of Ring fibrils. The percentages of segments corresponding to Ring-fold fibrils, straight filaments and non-aligned particles are indicated. Scale bar, 5 nm. **c**, 2D projection along the helical axis of Ring fibrils with the Ring fold, generated from reference-free 2D class averages. Scale bar, 5 nm. **d**, XY cross-sectional view of the 3D reconstruction of Ring fibrils with the Ring fold. Helical axis is shown as cross. Scale bar, 10 Å. **e**, Representative cryo-EM micrograph of Dot fibrils. Scale bar, 50 nm. **f**, Representative reference-free 2D class averages of segments from the cryo-EM dataset of Dot fibrils. The percentages of segments corresponding to Dot I, Dot II and Dot III fibrils, straight I and straight II filaments, and non-aligned particles are indicated. Scale bar, 5 nm. **g**, 2D projection along the helical axis of Dot I, Dot II, and Dot III fibrils, generated from reference-free 2D class averages. Scale bar, 5 nm. **h**, XY cross-sectional view of the 3D reconstructions of Dot Ia, Dot Ib and Dot II fibrils. Helical axis is shown as cross. Scale bar, 10 Å.

**Extended Data Fig. 4.**
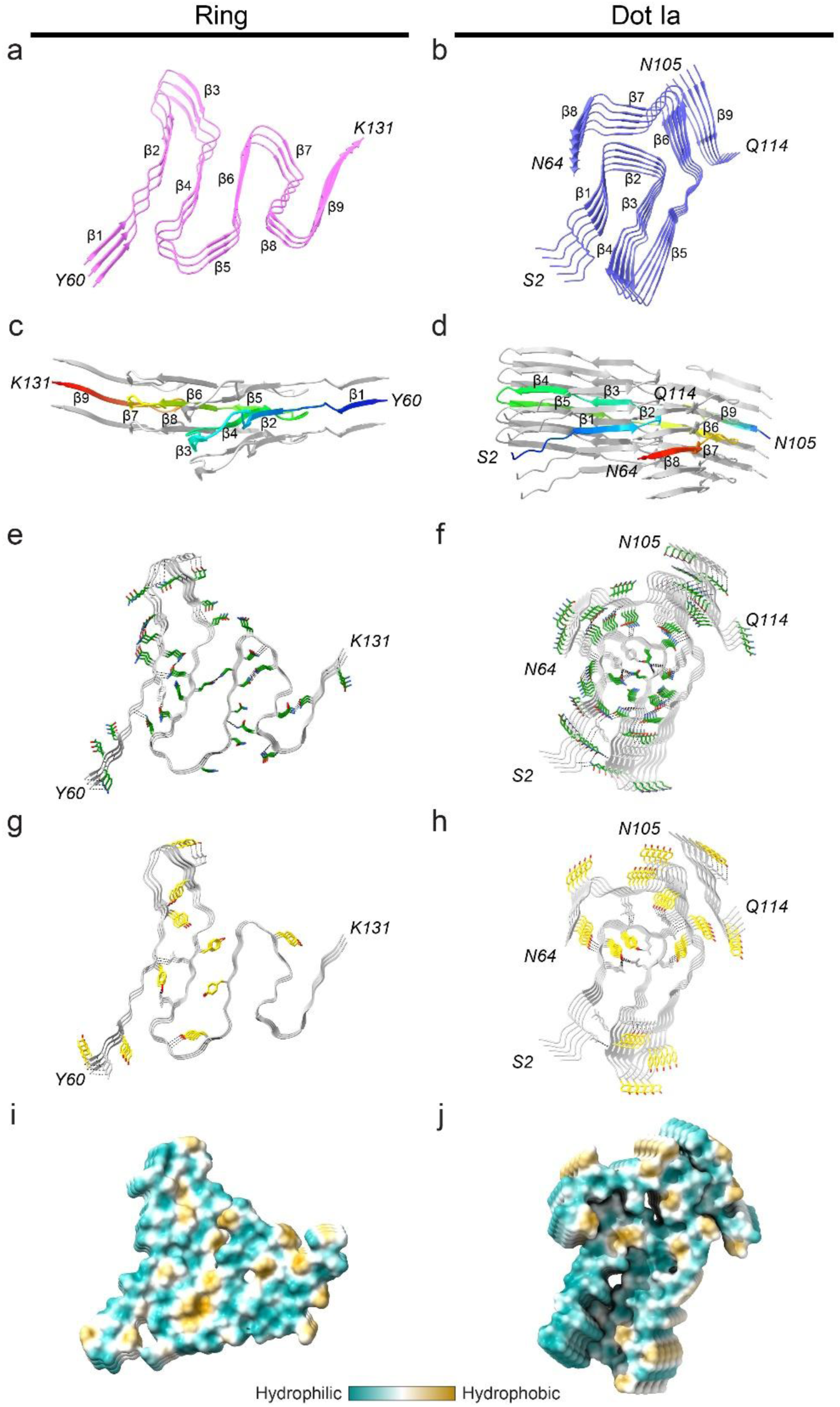
The amyloid structures of Sup35 Ring fold and Dot Ia fold. **a**, **b**, Structures of Ring fold and Dot Ia fold, shown with secondary-structure elements for three and five Sup35 molecules, respectively. **c, d**, Side views of the Ring fold and Dot Ia fold aligned with the helical axis, with the central layer coloured in rainbow to indicate height variation within a single layer. **e-h**, Views of the Ring fold and Dot Ia fold highlighting glutamine and asparagine residues (green; **e, f**), tyrosine residues (yellow; **g, h**) and their hydrogen bonding network (dashed lines). **i, j**, Hydrophobicity of the Ring fold and Dot Ia fold, ranging from most hydrophilic (cyan) to most hydrophobic (yellow).

**Extended Data Fig. 5.**
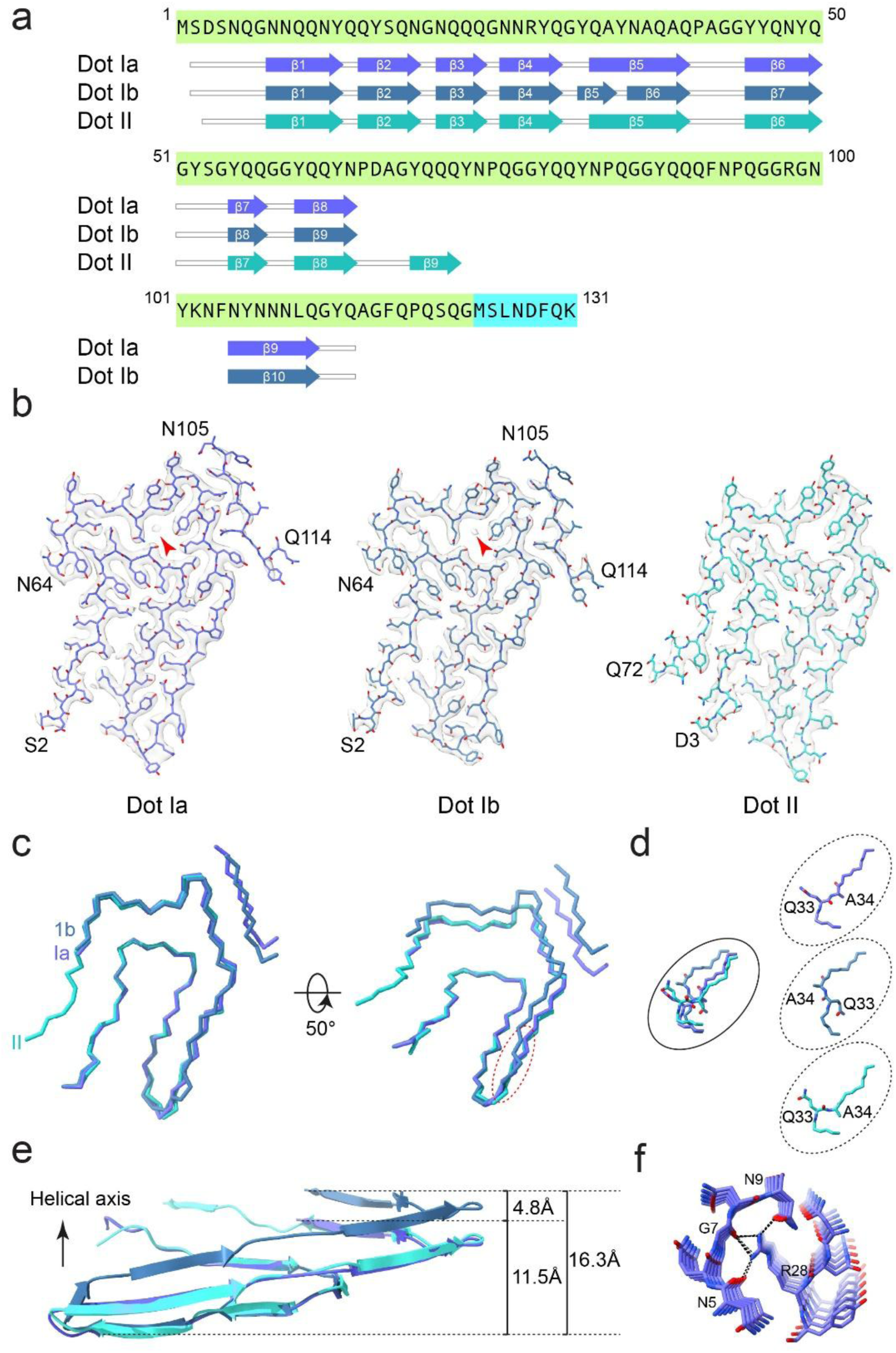
Structures of the Sup35 Dot Ia, Dot Ib and Dot II folds. **a**, Sequence alignment of secondary-structure elements in the Sup35 Dot Ia, Dot Ib and Dot II amyloid folds. Arrows indicate β-strands. **b**, Cryo-EM density maps, shown in transparent grey with fitted atomic models, of the Dot Ia (blue), Dot Ib (dark blue) and Dot II (blue-green) folds. **c**, Backbone comparison of Dot Ia (blue), Dot Ib (dark blue) and Dot II (blue-green), aligned over residues 3-30. **d**, Close-up views of the region around residues Q33 and A34. **e**, Comparison of secondary-structure elements in the Sup35 Dot fibril structures viewed perpendicular to the helical axis. **f**, Close-up view of structure around R28 in Dot Ia fold. Hydrogen bonds to the side chain of R28 are shown as dashed black lines.

**Extended Data Fig. 6.**
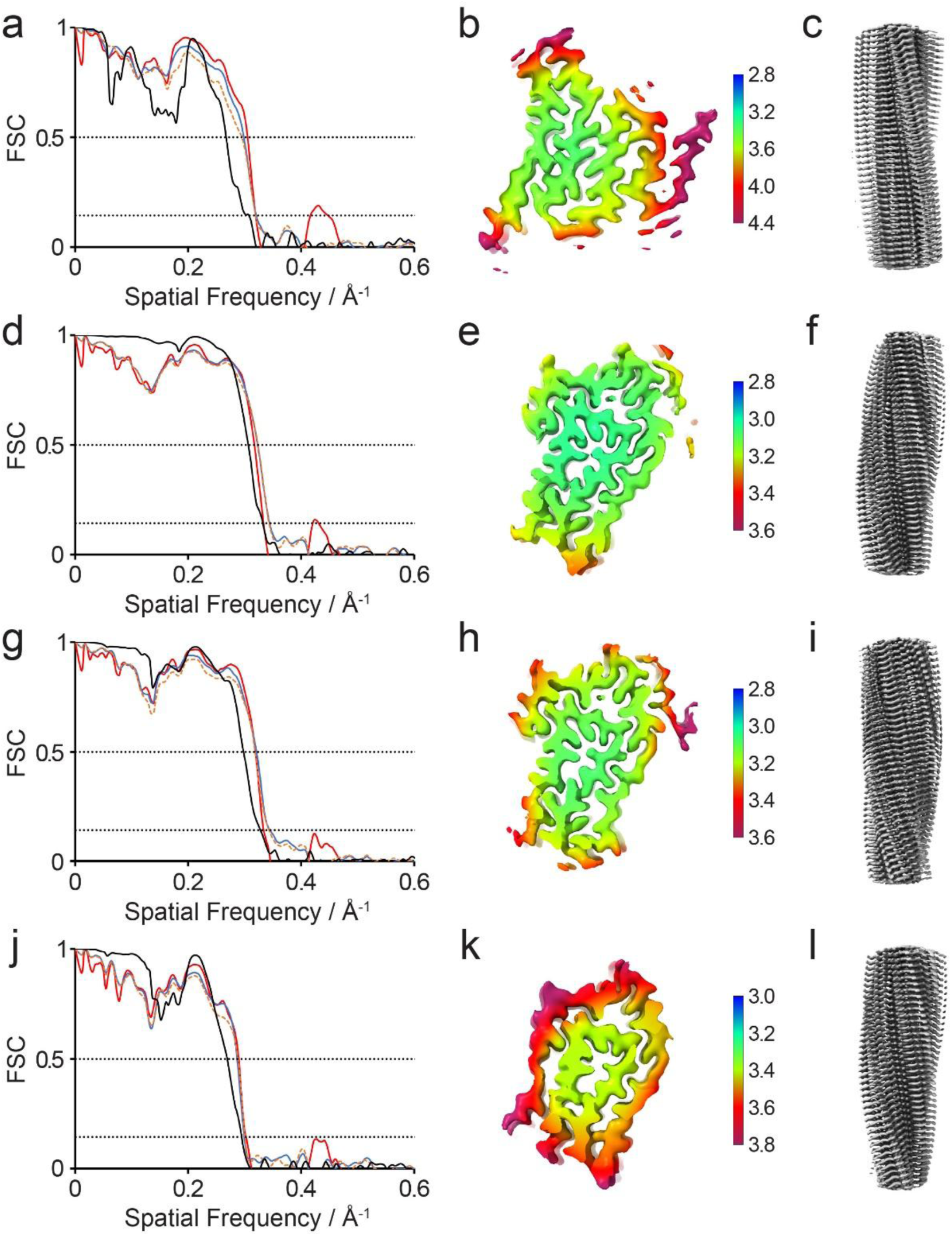
FSC curves and 3D reconstructions of Sup35 Ring and Dot fibrils. Fourier shell correlation (FSC) curves between two independently refined half-maps (black, solid), between the final model and the full map (red, solid), between a model refined against the first half-map and the first half-map (blue, solid), and between the same model and the second half-map (yellow, dashed) for Ring (**a**), Dot Ia (**d**), Dot Ib (**g**) and Dot II (**j**) folds. Local-resolution estimates for the 3D reconstructions of Ring (**b**), Dot Ia (**e**), Dot Ib (**h**) and Dot II (**k**) folds. Maps viewed along the helical axis, showing well resolved individual Sup35 molecules in Ring (**c**), Dot Ia (**f**), Dot Ib (**i**) and Dot II (**l**) folds.

**Extended Data Fig. 7.**
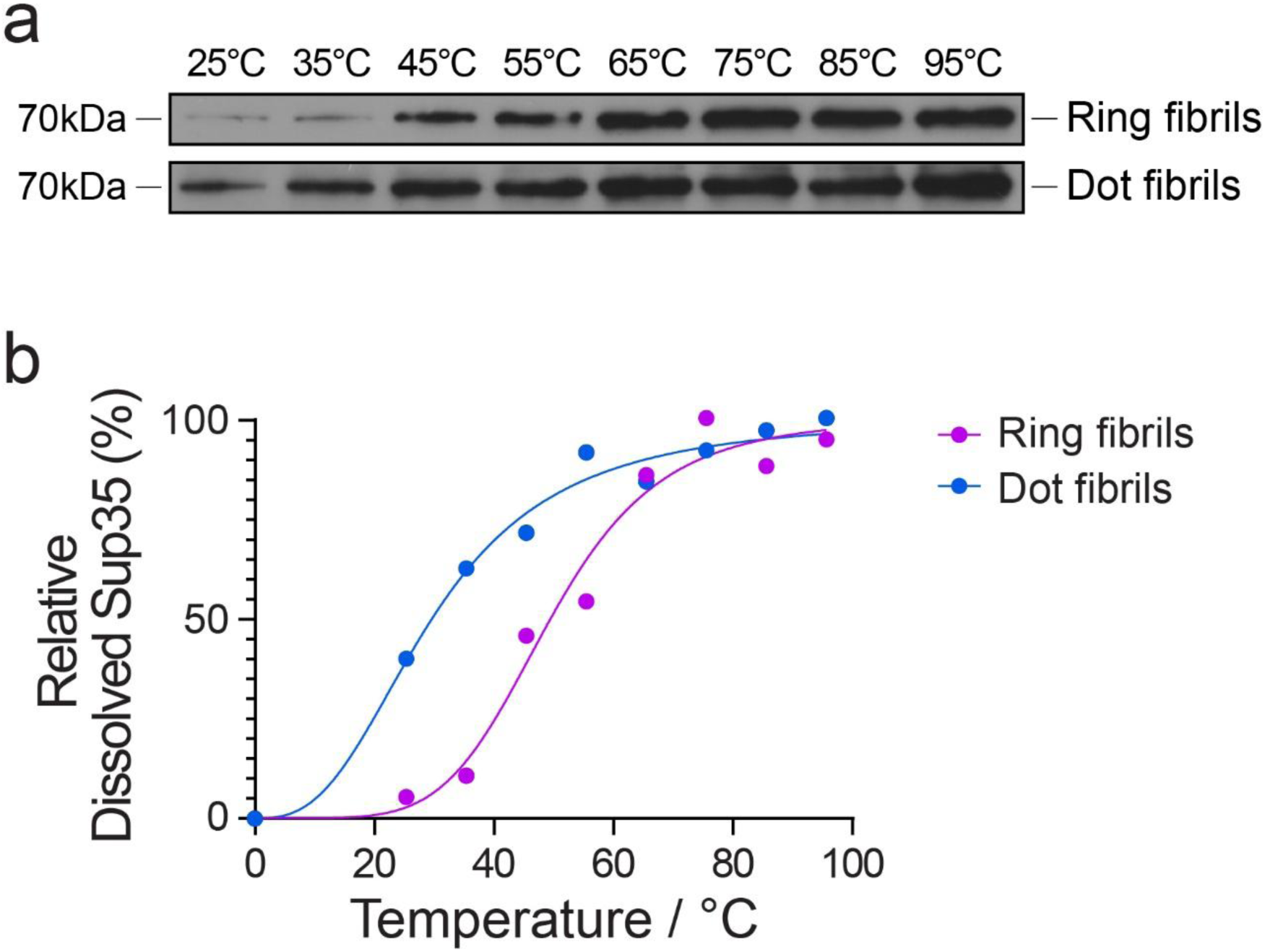
Thermostability assay of Sup35 Ring and Dot fibrils. **a**, SDS-PAGE analysis of the susceptibility of Ring and Dot fibrils to thermal solubilisation. **b**, Band intensities from **a** plotted against temperature, with curves fitted to a sigmoidal function.

**Extended Data Fig. 8.**
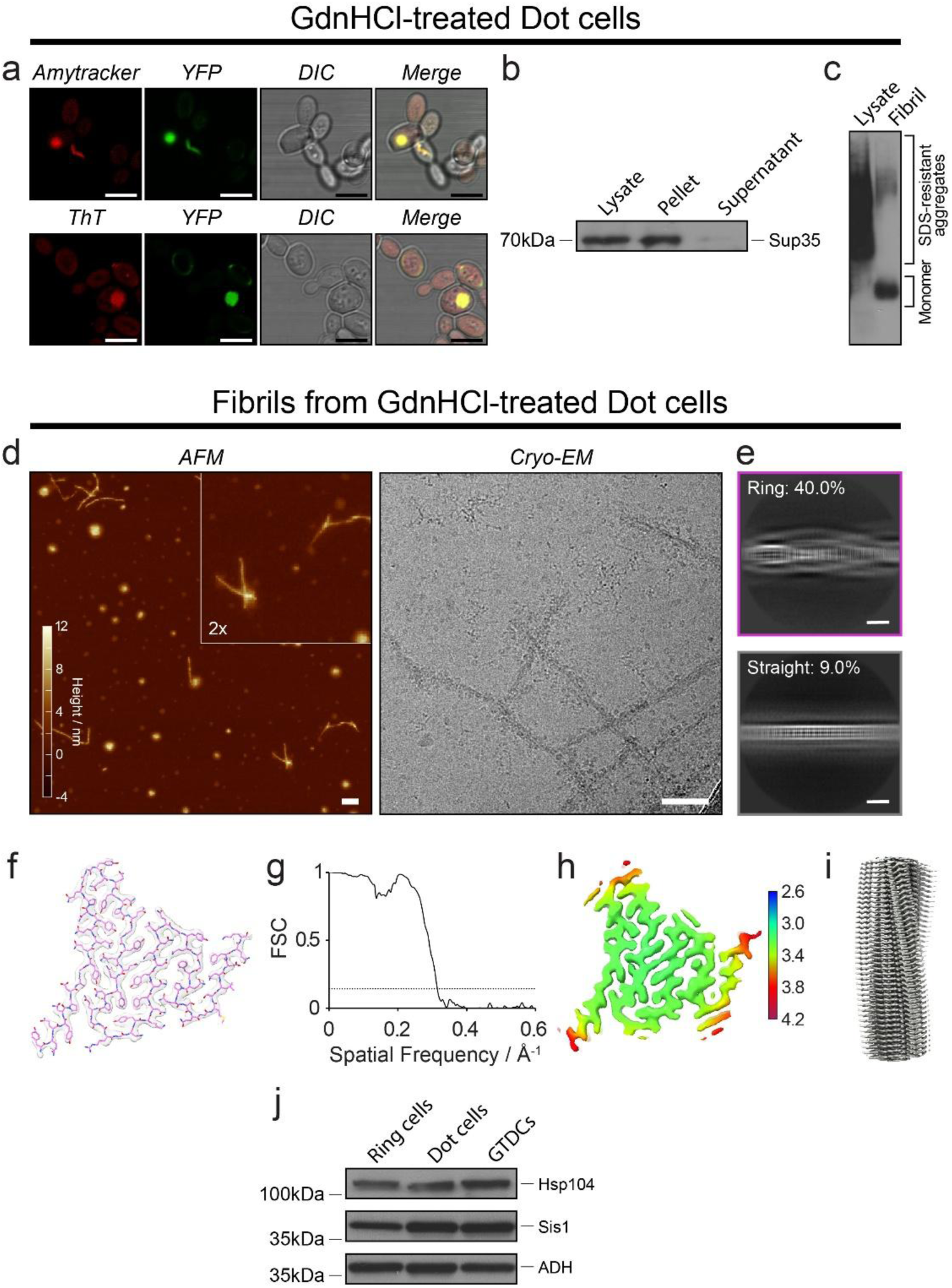
Characterisation of Sup35 fibrils purified from GdnHCl-treated Dot cells. **a**, Fluorescence microscopy images showing co-localisation of Amytracker and ThT with ring-like and dot-like aggregates in GdnHCl-treated Dot cells (GTDCs). Scale bar, 5µm. **b**, Sedimentation assay of GTDC lysate showing that Sup35 is predominantly aggregated. Antibody: anti-Sup35. **c**, SDD-AGE of GTDC fibrils followed by immunoblotting with anti-Sup35 antibody. **d**, AFM and Cryo-EM images of GTDC fibrils. Scale bar, 50 nm. **e**, Representative reference-free 2D class averages of segments from the cryo-EM dataset of GTDC fibrils. Scale bar, 5nm. **f**, Cryo-EM density map (transparent grey) with the Ring fold atomic model (pink) rigid-body fitted. **g**, FSC curves between two independently refined half-maps. **h**, Local-resolution estimates of the 3D reconstruction of GTDC fibrils. **i**, Map viewed along the helical axis, showing well-resolved individual Sup35 molecules. **j**, Western blot analysis of Hsp104 and Sis1 expression in lysates from Ring cells, Dot cells and GTDCs. Alcohol dehydrogenase (ADH) was used as a loading control.

**Extended Data Fig. 9.**
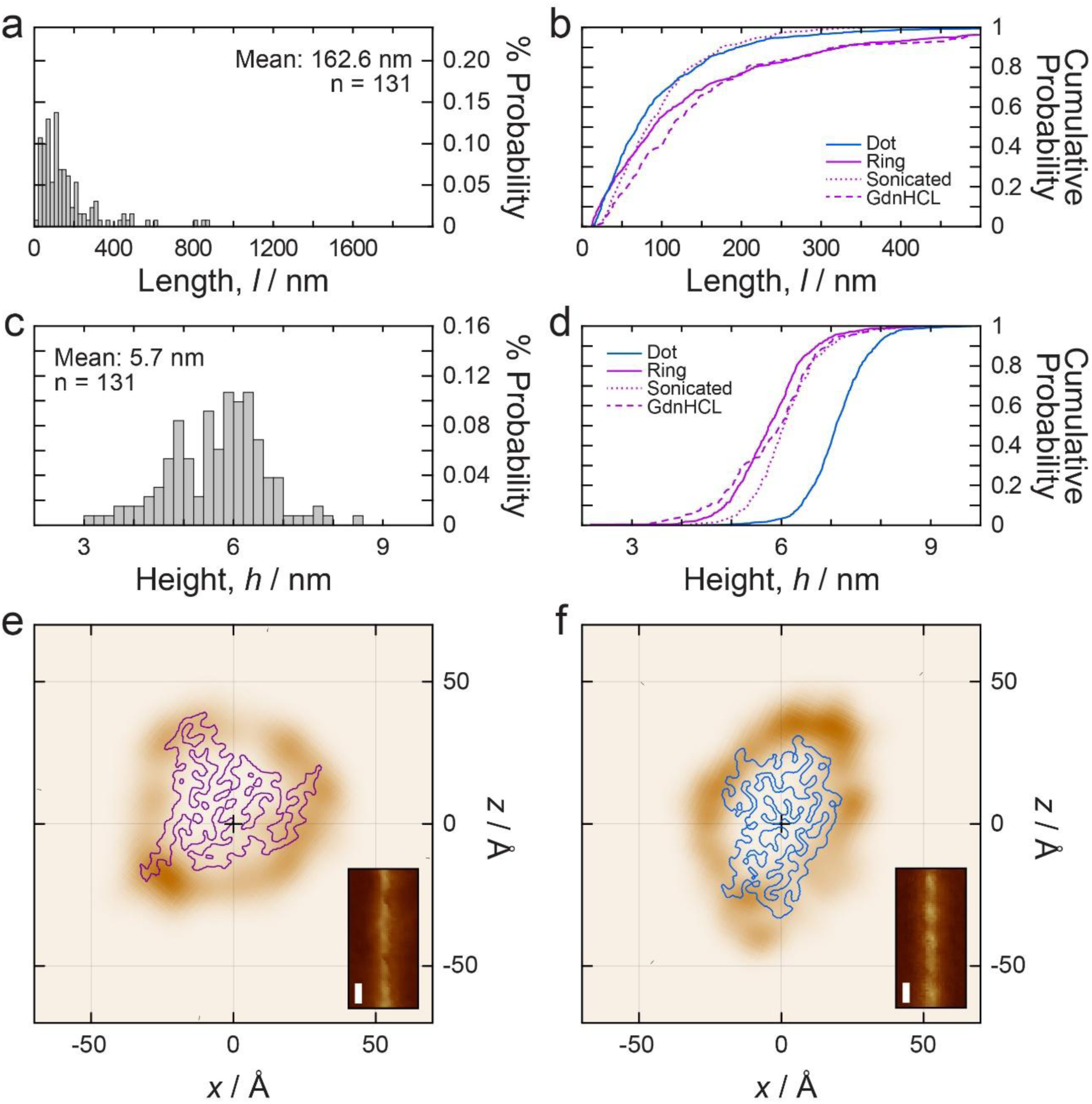
GdnHCl-treated Dot cell fibril population resembles the Ring state. **a**, Length distribution of Sup35 GTDC fibrils. **b**, Comparison of the length distribution of GTDC fibrils with Ring, Dot and Sonicated Ring fibrils. **c**, Height distribution of Sup35 GTDC fibrils. **d**, Comparison of the height distribution of GTDC fibrils with Ring, Dot and Sonicated Ring fibrils. **e**, **f**, Cross-sectional CPR-AFM density maps of individual fibrils from the GTDC fibril population that best matched the Ring (**e**) and Dot Ia (**f**) structures. The darker brown regions in these cross-sectional density maps indicate areas of contact between the AFM probe tip and the filament cross-section, and are shown in their best-fit orientations together with outlines of the corresponding cross-sections from the cryo-EM density maps. Insets show the corresponding digitally straightened image data.

**Extended Data Fig. 10.**
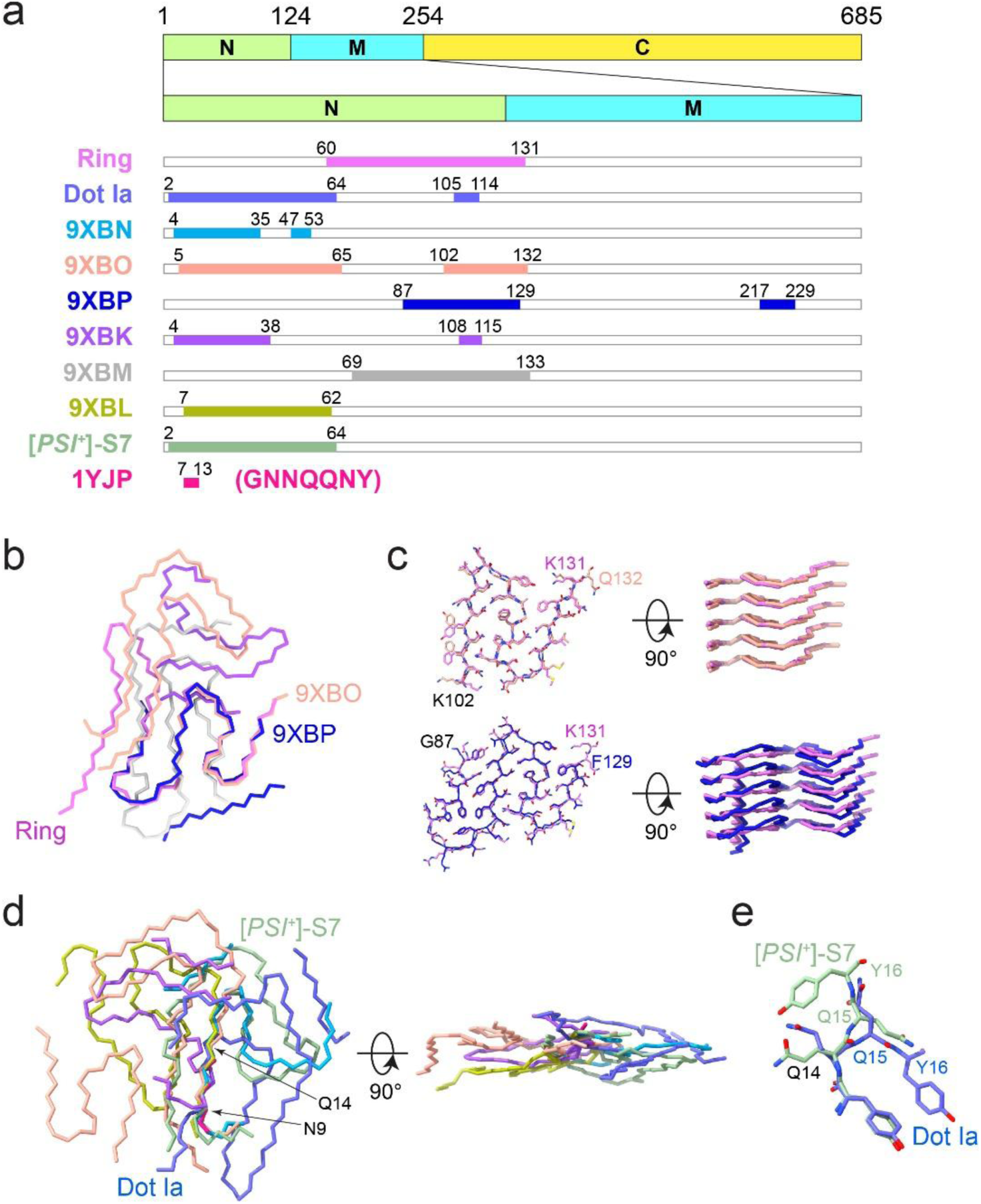
Structural comparison of Sup35 fibrils. **a**, Residue ranges of the ordered cores of the Ring and Dot Ia folds aligned with those of previously reported Sup35 fibril structures. The colours of named structures are used consistently for identification from **a-e**. **b**, Structural comparison of the Ring fold with other Sup35 amyloid structures containing overlapping ordered-core residue ranges. Because no single common residue range is shared across all structures, each structure was aligned to the Ring fold using its respective overlapping segment. **c**, Close-up comparison of the Ring fold with fibrils formed by recombinant wild-type Sup35 at 37 °C (Sc37, 9XBO) and by the S17R mutant at 4 °C (S17R4C, 9XBP). **d**, Structural comparison of the Dot Ia fold with other Sup35 amyloid structures containing overlapping ordered-core residue ranges. Residues 9-14 were used for alignment, except 1YJP, for which residues 9-13 were used. **e**, Close-up view showing that a 180° rotation of the peptide bond between Q15 and Y16 distinguishes the Dot Ia fold from the *ex vivo* [*PSI^+^*]-S7 structure and all other reported Sup35 fibril structures, although the Dot Ia fold, excluding the peptide island, spans the same residue range as the core of [PSI+]-S7 fibrils.

